# Cold and warmth intensify pain-linked sodium channel gating effects and persistent currents

**DOI:** 10.1101/2022.12.19.521077

**Authors:** Sophia Kriegeskorte, Raya Bott, Martin Hampl, Alon Korngreen, Ralf Hausmann, Angelika Lampert

## Abstract

Voltage-gated sodium channels (Na_v_) are key players in excitable tissues with the capability to generate and propagate action potentials. Mutations in the genes encoding Na_v_s can lead to severe inherited diseases, and some of these so-called channelopathies are showing temperature sensitive phenotypes, for example paramyotonia congenita, Brugada-syndrome, febrile seizure syndromes and inherited pain syndromes like erythromelalgia (IEM) and paroxysmal extreme pain disorder (PEPD). Nevertheless, most investigations of mutation-induced gating effects were conducted at room temperature and thus the role of cooling or warming in channelopathies remains poorly understood. Here, we investigated the temperature sensitivity of four Na_v_ subtypes: Na_v_1.3, Na_v_1.5, Na_v_1.6, and Na_v_1.7 and two mutations in Na_v_1.7 causing IEM (Na_v_1.7/L823R) and PEPD (Na_v_1.7/I1461T), using an automated patch clamp system. Our experiments at 15 °C, 25 °C and 35 °C revealed a shift of the voltage dependence of activation to more hyperpolarized potentials with increasing temperature for all investigated subtypes. Na_v_1.3 exhibited strongly slowed inactivation kinetics compared to the other subtypes that resulted in enhanced persistent current especially at 15 °C, indicating a possible role in cold induced hyperexcitability. Impaired fast inactivation of Na_v_1.7/I1461T was significantly enhanced by cooling temperature to 15 °C. The subtype specific modulation as well as the intensified mutation induced gating changes stress the importance to consider temperature as regulator for channel gating and its impact on cellular excitability as well as disease phenotypes.

**Summary:** Activation of the sodium channel subtypes Na_v_1.3, Na_v_1.5, Na_v_1.6, and Na_v_1.7 and two pain linked mutations is alleviated by warmth. Cooler temperatures, on the other hand, strongly enhance persistent currents of Na_v_1.3. The impaired fast inactivation of the pain-linked Na_v_1.7/I1461T mutation is further impaired by cooling, mimicking clinical findings.

## Introduction

Voltage-gated sodium channels (Na_v_) play an essential role in the electrical signaling of cells. With their capability to activate and inactivate rapidly in response to changes in membrane voltage they are initiating the upstroke of action potentials and are key players in excitable tissues. Nine Na_v_ α-subunits, Na_v_1.1-1.9, and four auxiliary β-subunits have been identified in humans so far (Catterall et al., 2005, Catterall, 2000, Morgan et al., 2000, Yu et al., 2003). Na_v_1.1, Na_v_1.2, Na_v_1.3, and Na_v_1.6 are mainly, but not exclusively, expressed in the central nervous system (CNS) (Liang et al., 2021, Whitaker et al., 2000, Vacher et al., 2008). Na_v_1.4 is mainly expressed in skeletal and Na_v_1.5 in cardiac muscles (Rogart et al., 1989, Trimmer et al., 1989), while Na_v_1.7, Na_v_1.8 and Na_v_1.9 are found in the peripheral nervous system (PNS) (Bennett et al., 2019, Fukuoka et al., 2008).

The voltage dependence of activation and inactivation, their time constants as well as the recovery from inactivation are, like all cellular processes, influenced by temperature changes. Nevertheless, comprehensive studies characterizing temperature dependent gating are rare. Because Na_v_s are an important factor for the overall excitability of neurons, serious functional consequences are related to temperature induced changes in their gating properties. For example, enhanced functionality of Na_v_1.2 at febrile (40 °C to 41 °C) compared to physiological (36 °C) temperature mediated the increase in neuronal excitability in *in-vitro* experiments with cortical tissue (Ye et al., 2018). Computer simulations revealed that already small elevations in temperature from physiological to fever conditions increase the excitability of central neurons, expressed in higher firing rates and faster action potential conduction velocity (Ye et al., 2018). *In-vivo* experiments with mice, which were exposed to a 42 °C environment for 30 min, showed that febrile temperature alone can provoke seizure-related behavioral changes (Ye et al., 2018). Moreover, gain or loss of function mutations in Na_v_s cause severe, temperature provoked diseases, like Brugada syndrome (Samani et al., 2009, Keller et al., 2005), Paramyotonia congenita (PMC) (Bouhours et al., 2004, Carle et al., 2009, Ke et al., 2017), febrile epileptic syndromes (Volkers et al., 2013, Peters et al., 2016) and inherited pain syndromes (Körner and Lampert, 2020).

Na_v_s expressed in peripheral nerve endings of the skin are necessary to generate and propagate action potentials encoding sensory information. Compared to Na_v_s of the CNS, they are exposed to much larger variations in temperature. The tetrodotoxin (TTX)-resistant Na_v_1.8 is necessary to transduce nociceptive information in sensory neurons at low temperature in mice (Zimmermann et al., 2007). While at 10 °C TTX-sensitive Na_v_s are mostly slow inactivated at the resting membrane potential, Na_v_1.8 remains excitable. To encode noxious heat, the TTX-resistant Na_v_1.9 is required in rodents and undergoes a large gain of function with increasing temperature (Touska et al., 2018). Thus, it is likely that different Na_v_ isoforms have different intrinsic sensitivity to temperature changes.

Rare inherited chronic pain syndromes with temperature sensitive phenotypes can be caused by mutations in Na_v_1.7 (Körner and Lampert, 2020). Inherited erythromelalgia (IEM), which is characterized by burning pain, redness and warmth of the extremities, can be provoked by elevated ambient temperature and exercise (van Genderen et al., 1993). In contrast, mutations causing paroxysmal extreme pain disorder (PEPD) lead to attacks of rapidly developing burning pain in regions of the rectum, eye and mandibular, that may be accompanied by vegetative symptoms (Bennett et al., 2019). In this pain syndrome cooling, for example cold wind, is described as one possible trigger factor (Fertleman et al., 2007). Most of the known IEM mutations cause shifts in the voltage dependence of half-maximal (V_1/2_) activation to more hyperpolarized potentials, while the PEPD mutations mainly impair steady-state fast inactivation (Baker and Nassar, 2020, Tang et al., 2015, Jarecki et al., 2008). The effects observed have been linked to neuronal hyperexcitability causing pain. Here, we focus exemplary on the IEM linked mutation Na_v_1.7/L823R, that inserts an additional positive charge in the voltage sensor of Domain II (DII) (Lampert et al., 2009), and the PEPD mutation Na_v_1.7/I1461T, changing the unipolar isoleucine of the inactivation-motif to a polar threonine (Fertleman et al., 2006).

Neuropathic pain, for example after a nerve or spinal cord injury, is often accompanied by cold allodynia, in which already innocuous cold stimuli lead to pain (Jensen and Finnerup, 2014). In the last decades, a lot of effort has been put into the investigation of sensory transducers that are responsible for the detection of temperature. Thermosensitive transient receptor potential ion channels (TRPs) have been identified to enable somatosensory neurons to sense temperature (Caterina et al., 1997, Tominaga et al., 1998), with TRPM8 being sensitive to cool temperatures (McKemy, 2005, McKemy et al., 2002). Interestingly, it was shown in mouse models that after chronic constriction injury of a nerve leading to chronic pain the number of TRPM8 expressing dorsal root ganglia (DRG) neurons as well as their functional properties are unaltered. Still, TRPM8 is required for cold allodynia, which is absent in TRPM8 KO mice (Caspani et al., 2009, Knowlton et al., 2013, Colburn et al., 2007). Previously, silent cold-sensing neurons have been described to contribute to cold allodynia in neuropathic pain states and K_v_1 potassium channels were identified as potential cold sensors (MacDonald et al., 2021). There is also evidence that Na_v_s are crucial for cold induced pain in different neuropathic pain conditions (Sittl et al., 2012, Zimmermann et al., 2013).

Despite the temperature sensitivity of several Na_v_-channelopathies, information about temperature induced changes of gating properties is mostly lacking. Most electrophysiological experiments are, due to the technical challenges at physiological temperature, conducted at room temperature and reliable data of electrophysiological characterizations at different temperatures is rare. A few studies have investigated the effect of temperature on different Na_v_ subtypes. Most of them revealed that with increasing temperature the speed of gating, reflected in the time constants, is accelerated, while the effect on the voltage dependence of steady-state activation and inactivation is still under debate and varies depending on the experimental conditions (Ruff, 1999, Sarria et al., 2012, Thomas et al., 2009, Ye et al., 2018, Zimmermann et al., 2007, Egri et al., 2012).

In this study, we investigated four different Na_v_ subtypes, Na_v_1.3, Na_v_1.5, Na_v_1.6, and Na_v_1.7, regarding their temperature dependent gating. We studied two mutations of Na_v_1.7, causing IEM and PEPD, in order to test whether the clinical phenotype is reflected in their gating properties. We performed high-throughput patch clamp experiments with an automated patch clamp device, the SyncroPatch 384, at 15 °C, 25 °C and 35 °C, which revealed a subtype-specific modulation by temperature as well as intensified mutation induced gating changes. The voltage dependence of activation was shifted to more hyperpolarized potentials with increasing temperature for all investigated subtypes, with an enhanced left shift in the IEM mutation, while V_1/2_ of steady-state fast inactivation was in general less affected. Our results stress the importance to consider temperature as a regulator for channel gating and its impact on cellular excitability and disease phenotype.

## Materials and methods

### Cell culture and cell preparation

All investigated Na_v_ subtypes (rat (r) Na_v_1.3, human (h) Na_v_1.5, mouse (m) Na_v_1.6, hNa_v_1.7, hNa_v_1.7/L823R, and hNa_v_1.7/I1461T) were stably expressed in cells from the human embryonic kidney cell line HEK293 and kept under standard culture conditions (37 °C and 5 % CO_2_). Cell culture media and supplements are listed in the supplementary data S1, Table S1.

All electrophysiological experiments were performed with cells at passage numbers lower 30 and after reaching confluency of 70 – 95 %. One day prior to the recording the media was refreshed and the cells were incubated at 32 °C and 5 % CO_2_.

In order to carry out the experiments, the cells were washed twice with PBS-EDTA (PBS: PAN-Biotech, Aidenbach, Germany; EDTA: Sigma-Aldrich, St. Louis, Missouri, USA), then treated with Trypsin/EDTA (Sigma-Aldrich, St. Louis, Missouri, USA) and incubated at 37 °C for 6 min to detach, before “External –Mg^2+^ –Ca^2+^” (Nanion Technologies GmbH, Munich, Germany) containing (in mM): 10 HEPES, 140 NaCl, 5 Glucose and 4 KCl was added. Subsequently, the cells were stored at 4 °C for 10 min. Cells were pipetted up and down 8 - 10 times using fire-polished glass pipettes to break up cell clumps, “External –Mg^2+^ –Ca^2+^” was added until a final volume of 30 ml and cells were transferred to the “Cell Hotel” on the SyncroPatch 384 (Nanion Technologies GmbH, Munich, Germany), where they rested at least 30 min at 10 °C and shaking speed of 333 RPM before the recording was started.

### Electrophysiology

Whole-cell voltage clamp recordings at 15 °C, 25 °C and 35 °C were performed using the high-throughput patch clamp robot SyncroPatch 384 with “NPC-384T 1x S-Type” chips (2 µm holes, Nanion Technologies GmbH, Munich, Germany) and, for data acquisition, PatchControl384 Version 1.9.7 (Nanion Technologies GmbH, Munich, Germany). Whole-cell recordings were conducted according to Nanion’s procedure, including initialization, cell-catch, sealing, whole-cell formation, liquid application and recordings. After achieving whole cell configuration, capacitive transients were cancelled, series resistance compensation was set to 65 %, leak currents were subtracted online using a P/4 procedure and signals were sampled at 20 kHz. The median series resistance was always lower than 6.1 MΩ.

The integrated temperature control unit of the SyncroPatch 384 allowed the adjustment of the temperature in a range from 10 °C to 43 °C throughout the experiment. The temperature of both, solution and measurement chamber, was set to either 15 °C, 25 °C or 35 °C. Experiments at each temperature step were performed successively with different cells from one batch.

The intracellular solution “Internal CsF110” contained (in mM): 10 EGTA, 10 Hepes, 10 CsCl, 10 NaCl and 110 CsF. The extracellular solution “External Standard” (only used for recordings of Na_v_1.6) contained (in mM): 140 NaCl, 10 Hepes, 5 Glucose, 4 KCl, 2 CaCl_2_ and 1 MgCl_2_, while the extracellular solution “External NMDG 60” (used for the recordings of all other channels) contained (in mM): 80 NaCl, 60 NMDG, 10 Hepes, 5 Glucose, 4 KCl, 2 CaCl_2_ and 1 MgCl_2_ (all solutions made by Nanion Technologies GmbH, Munich, Germany).

The channels activation was assessed using 30 ms pulses to a range of test potentials (−85 mV to + 30 mV) from the resting membrane potential of -120 mV in 5 mV steps with an interval of 5 s at resting membrane potential between each sweep. The conductance-voltage (G-V) relationship, the inactivation time constant *τ* as well as the persistent currents were determined from these recordings. The voltage dependent Na_v_ conductance *G*_*Na*_ was calculated according to

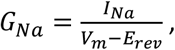

where *I*_*Na*_ is the peak current at the voltage *V*_*m*_ and *E*_*rev*_ is the reversal potential for sodium, determined for each cell individually. Activation curves were derived by plotting *G*_Na_ normalized to the maximum conductance *G*_*Na,max*_ as a function of test potential and fitting it with the Boltzmann distribution equation:

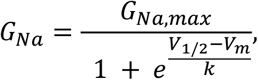

where *V*_*1/2*_ is the membrane potential at half-maximal activation, *V*_*m*_ is the membrane voltage and *k* is the slope factor. Persistent sodium current *I*_*pers*_ was defined as the mean remaining current between 26.5 to 29.5 ms for each 30 ms activation pulse and normalized to the peak current *I*_*peak*_. It is plotted as a function of test potential. The population average of all included traces was used to determine the inactivation time constant *τ* with a mono-exponential fit on the decay part of this traces. To quantitatively determine the dependence of the inactivation process on temperature, it was described by the Arrhenius equation:

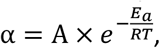

where *α* is the forward rate constant in the transition between open and inactivated state, *O ⇆ I*, A is a proportionality constant, *E*_*a*_ is the activation energy, *R* is the gas constant and *T* is the absolute temperature. Since *τ ≈ 1/α* at depolarized potentials, it was possible to use the Arrhenius plot to estimate *E*_*a*_ by plotting ln(τ) as a function of *1/T*, because the slope corresponds to *E*_*a*_*/R* in this case. At more negative potentials *τ = 1/(α+β)* and therefore the simple Arrhenius analysis is generally not applicable. The exception to this rule is when the activation energy for *α* and the backward rate constant *β* is the same allowing accurate calculation of *E*_*a*_ from the Arrhenius plots.

The voltage dependence of steady-state fast inactivation was measured using a series of 500 ms pre-pulses from -130 mV to -20 mV in 10 mV steps followed by a 40 ms test pulse to 0 mV that assessed the non-inactivated transient current. The normalized peak inward currents were fitted using a Boltzmann distribution equation:

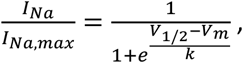

where *I*_*Na,max*_ is the peak sodium current elicited after the most hyperpolarized pre-pulse, *V*_*m*_ is the preconditioning pulse potential, *V*_*1/2*_ is the half-maximal sodium current and *k* is the slope factor.

The recovery from fast inactivation was measured using a 500 ms pre-pulse to 0 mV followed by a hyperpolarizing recovery-pulse to -100 mV of varying duration (1 ms – 2000 ms). Due to the amplifier settings, it was not possible to measure time intervals shorter than 1 ms. After that, another depolarizing test pulse to 0 mV was applied to assess the rate of recovered channels. The maximum inward current of the test pulse *I*_*recov*_ was normalized to the maximum inward current of the pre-pulse *I*_*pre*_ and plotted against the duration of the recovery-pulse. The following double-exponential equation was used:

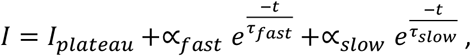

where *I* is the current amplitude, *I*_*plateau*_ is the amplitude at recovery time t=1ms, *α*_*fast*_ and *α*_*slow*_ are the amplitudes for time constants *τ*_*fas*t_ and *τ*_*slow*_ and *t* is time. *%*_*fast*_ is the fraction of the overall recovery that is accounted for by the faster recovering component following the equation:

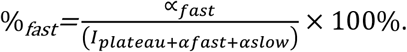

To evoke ramp currents, slow depolarizing pulses were applied from a holding potential of -120 mV to +5 mV or to +20 mV. Rates of 1.4 mV/ms, 2.5 mV/ms and 5 mV/ms were investigated and the maximum inward current *I*_*ramp*_ was normalized to the maximum inward current elicited in the G-V-relationship *I*_*Act*_.

5 ms voltage pulses to 0 mV from the holding potential of -120 mV were applied at frequencies of 20 Hz, 50 Hz and 100 Hz to assess the use-dependent current decay. The peak inward current of the 10^th^ action potential *I*_*10th*_ was normalized to the peak inward current of the first action potential *I*_*1st*_.

### Data analysis and statistics

The recorded data were analyzed using DataControl384 version 2.0.0 (Nanion Technologies GmbH, Munich, Germany), IgorPro (WaveMetrics, Lake Oswego, Oregon, USA) and Prism version 9 (GraphPad Software, San Diego, California, USA).

For statistical testing, groups of two were compared by Student’s t-test or a Mann-Whitney test in case of non-parametric testing. Groups larger than two were compared by an One-Way-Anova followed by Sidak’s multiple comparison test for parametric testing and Kruskal-Wallis test with Dunn’s multiple comparison in case of non-parametric testing. Statistical significance was defined as p < 0.005. All values are mentioned as mean ± SEM, unless otherwise stated..

### Supplemental material

Detailed information concerning the cell culture media and supplements for each cell line individually are listed in Table S1 in the supplementary information.

## Results

### Voltage dependence and kinetics of Na_v_ activation are modulated by temperature

The gating of Na_v_s is sensitive to temperature. Here, we investigated four wild-type isoforms: Na_v_1.3, Na_v_1.5, Na_v_1.6 and Na_v_1.7, each at 15 °C, 25 °C and 35 °C. Fig. 1 shows traces presenting the population average of all analyzed experiments.

**Figure 1.**
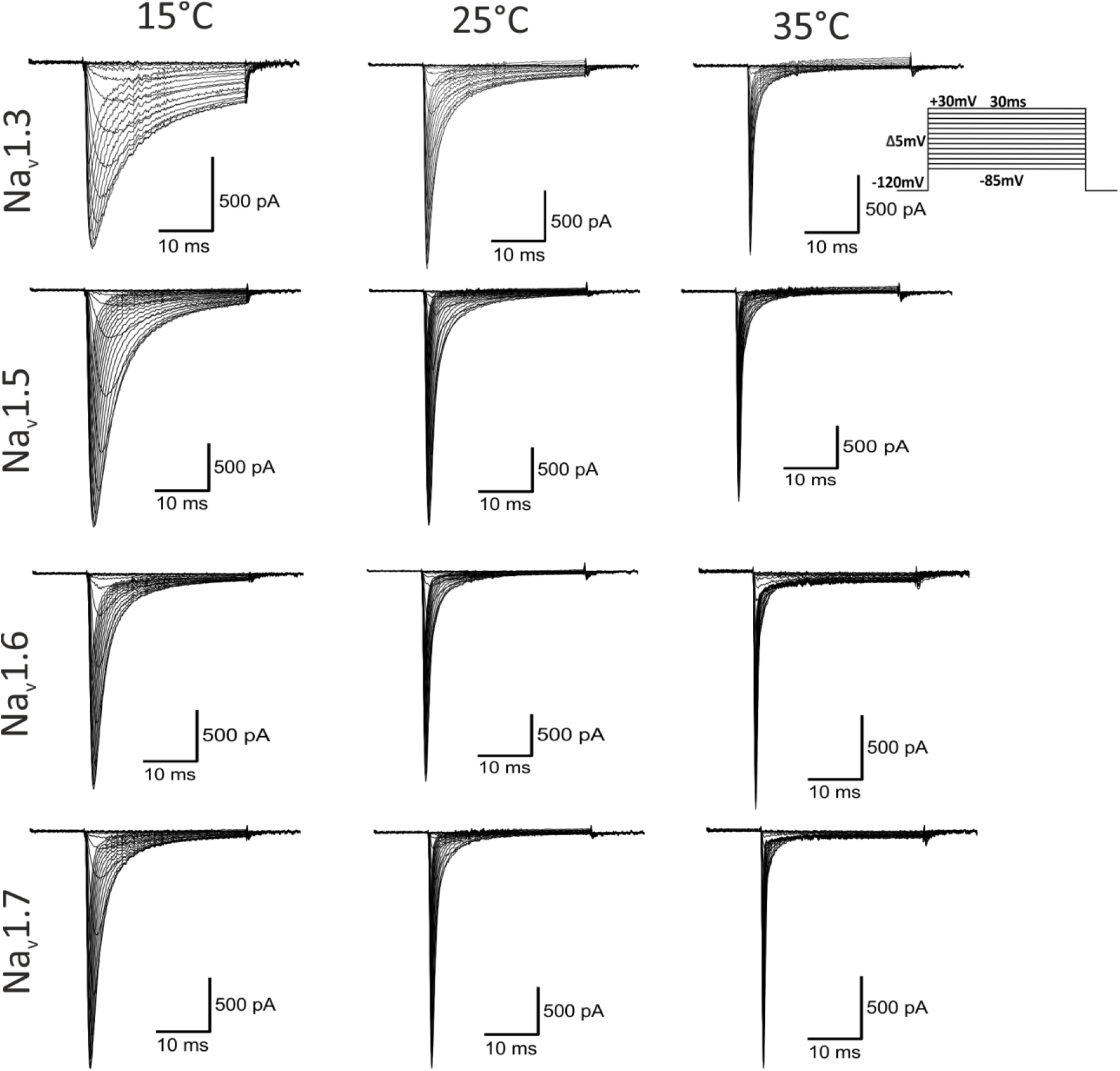
Population average of all analyzed experiments. Average of all included current traces, elicited by applying 30 ms - depolarizing pulses from a holding potential of -120 mV in 5 mV steps from -85 mV to 30 mV (voltage protocol in the inlet). Current traces of **(A)** Na_v_1.3, **(B)** Na_v_1.5, **(C)** Na_v_1.6 and **(D)** Na_v_1.7 at 15 °C, 25 °C and 35 °C. By presenting the average of all included recordings, a human selection bias that selects unusually nice recordings as representative traces was avoided.

First, we quantified the impact of temperature on Na_v_ activation. Without exception we found hyperpolarizing shifts of *V*_*1/2*_ in the range between 5.4 mV (Na_v_1.6) to 9.5 mV (Na_v_1.3) with increasing temperature from 15 °C to 35 °C (Fig. 2). These shifts were significant in both temperature increments for Na_v_1.3 (Fig. 2A) and Na_v_1.7 (Fig. 2D), while for Na_v_1.5 (Fig. 2B) and Na_v_1.6 (Fig. 2C) only the temperature rise from 15 °C to 25 °C produced a significant shift. Except for Na_v_1.6 at 35 °C, we also observed a steepening of the conductance curves and decreased slope factors *k* with warmer temperature (Fig. 2, Table 1).

**Figure 2.**
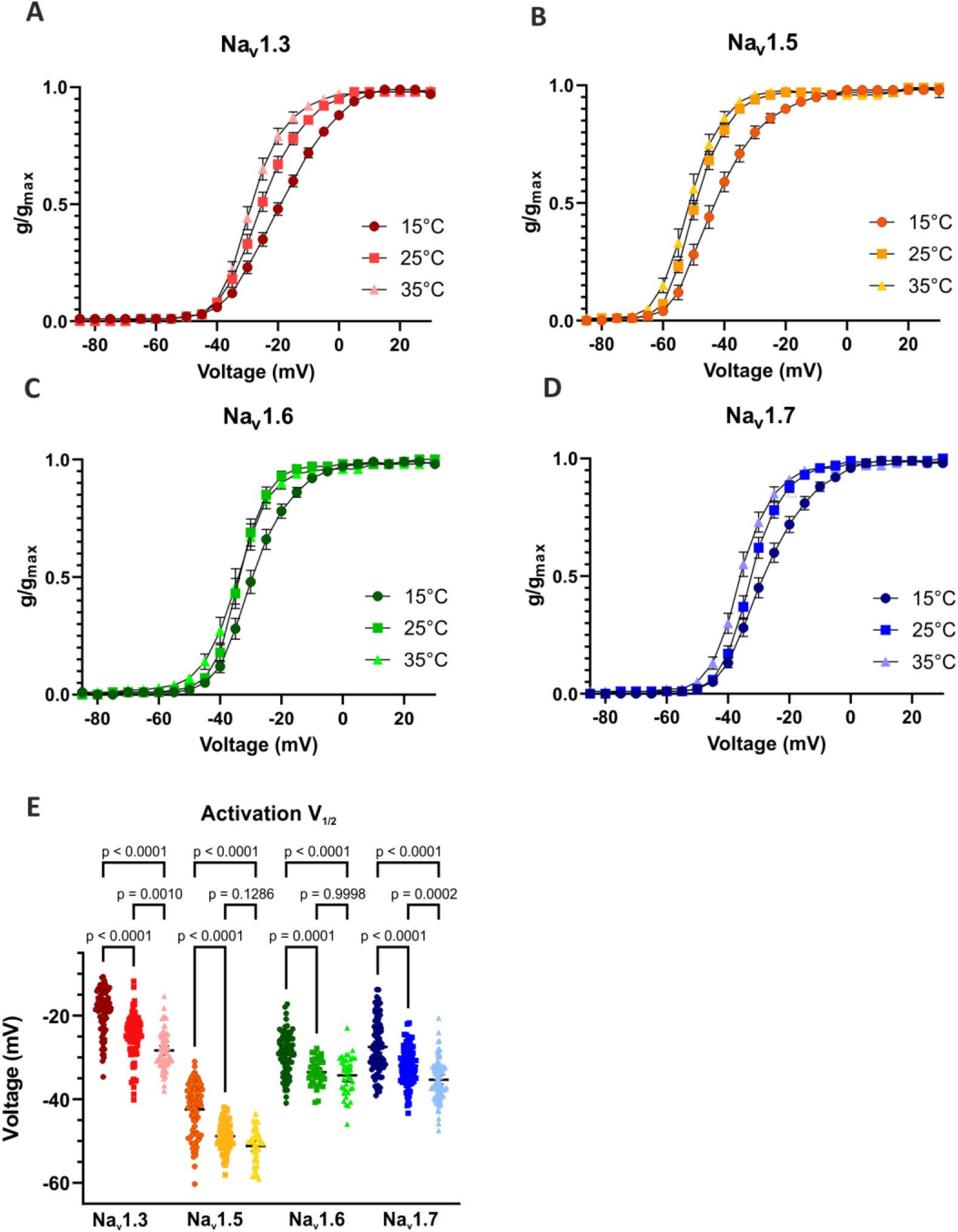
Warmer temperature shifts the voltage dependence of activation to more hyperpolarized potentials. The voltage dependence was shifted to more hyperpolarized potentials with warming from 15°C for all tested channel isoforms. G-V curves of **(A)** Na_v_1.3, **(B)** Na_v_1.5, **(C)** Na_v_1.6 and **(D)** Na_v_1.7. **(E)** Values of half-maximal voltage dependence of activation (V_1/2_) obtained from Boltzmann fits for individual traces. Means ± 95% confidence-interval. One-way ANOVA with Sidak’s multiple comparisons test.

**Table 1.** Summary of electrophysiological parameters determined for Na_v_1.3, Na_v_1.5, Na_v_1.6 and Na_v_1.7 (n.a.: not available)

|  | Nav1.3 |  |  | Nav1.5 |  |  | Nav1.6 |  |  | Nav1.7 |  |  |
| --- | --- | --- | --- | --- | --- | --- | --- | --- | --- | --- | --- | --- |
|  | 15°C | 25°C | 35°C | 15°C | 25°C | 35°C | 15°C | 25°C | 35°C | 15°C | 25°C | 35°C |
| **Activation parameter** |  |  |  |  |  |  |  |  |  |  |  |  |
| $V_{1/2}$ (mV) | -18.8 ± 0.5 | -25.0 ± 0.6 | -28.3 ± 0.6 | -42.5 ± 0.7 | -48.9 ± 0.4 | -51.2 ± 0.6 | -28.9 ± 0.6 | -33.5 ± 0.5 | -34.3 ± 0.8 | -27.5 ± 0.7 | -32.0 ± 0.5 | -35.4 ± 0.5 |
| $k$ | 8.2 ± 0.1 | 6.0 ± 0.2 | 4.7 ± 0.2 | 6.6 ± 0.2 | 4.7 ± 0.2 | 4.5 ± 0.2 | 5.3 ± 0.2 | 3.9 ± 0.2 | 4.6 ± 0.2 | 6.3 ± 0.2 | 4.5 ± 0.2 | 4.2 ± 0.2 |
| Time-to-peak (ms)<br>(0 mV) | 1.36 ± 0.05 | 0.53 ± 0.02 | n.a. | 0.78 ± 0.02 | 0.47 ± 0.01 | n.a. | 0.68 ± 0.01 | 0.45 ± 0.01 | n.a. | 0.64 ± 0.01 | 0.38 ± 0.005 | n.a. |
| $n$ | 90 | 84 | 64 | 80 | 77 | 50 | 71 | 36 | 38 | 94 | 80 | 121 |
| **Inactivation parameter** |  |  |  |  |  |  |  |  |  |  |  |  |
| $V_{1/2}$ (mV) | -68.6 ± 0.5 | -69.5 ± 0.7 | -67.4 ± 0.6 | -82.7 ± 0.6 | -82.4 ± 0.4 | -84.1 ± 0.8 | -65.9 ± 0.6 | -62.0 ± 0.5 | -64.2 ± 0.8 | -76.2 ± 0.9 | -75.4 ± 0.5 | -76.1 ± 0.7 |
| $k$ | -11.5 ± 0.2 | -9.2 ± 0.1 | -8.0 ± 0.1 | -8.3 ± 0.1 | -6.1 ± 0.1 | -5.6 ± 0.1 | -7.55 ± 0.1 | -6.0 ± 0.1 | -6.3 ± 0.1 | -10.3 ± 0.3 | -7.0 ± 0.1 | -7.3 ± 0.1 |
| $n$ | 93 | 102 | 99 | 94 | 128 | 96 | 71 | 78 | 74 | 138 | 135 | 121 |
| **Inactivation kinetics** |  |  |  |  |  |  |  |  |  |  |  |  |
| $\tau$ (ms) (0 mV) | 4.80 | 1.20 | 0.54 | 1.72 | 0.70 | 0.42 | 1.49 | 0.69 | 0.38 | 1.13 | 0.53 | 0.28 |
| Max $I_{pers}$ (% from $I_{max}$) | 21.0 ± 0.6 | 5.2 ± 0.2 | 1.1 ± 0.1 | 6.4 ± 0.3 | 2.0 ± 0.1 | 0.9 ± 0.1 | 3.6 ± 0.2 | 1.2 ± 0.1 | n.a. | 2.5 ± 0.3 | 0.7 ± 0.1 | n.a. |
| **Recovery from fast inactivation** |  |  |  |  |  |  |  |  |  |  |  |  |
| % $\tau$ fast | 76.3 | 69.9 | 37.1 | 71.0 | 70.1 | 62.3 | 81 | 75.3 | 39.5 | 87.0 | 81.4 | 51.0 |
| $\tau$ fast (ms) | 19.7 | 7.6 | 3.5 | 93.1 | 34.0 | 23.5 | 10.2 | 2.7 | 2.7 | 49.2 | 13.3 | 8.5 |
| $\tau$ slow (ms) | 250.9 | 285.5 | 193.4 | 570.0 | 287.4 | 246.8 | 398.6 | 303.9 | 252.3 | 447.8 | 428.8 | 621.9 |

We analyzed the time to peak, the time from onset of the voltage step until the maximum inward sodium current is reached. The channel opening was between 1.5 times (Na_v_1.6) and 2.6 times (Na_v_1.3) faster at 25 °C compared to 15 °C. At 35 °C, the activation became too fast to be accurately dissolved and no reliable analysis was possible. Taken together, the hyperpolarized *V*_*1/2*_ as well as the accelerated opening suggest a strong impact of temperature on the overall excitability of Na_v_s.

### Effects of temperature on the inactivation properties of Na_v_ subtypes

Na_v_s inactivate quickly within milliseconds upon activation. To assess the channels steady-state fast inactivation, we used the voltage protocol shown in Fig. 3A. No significant shifts in the voltage dependence of fast inactivation caused by temperature variation were observed, except for Na_v_1.6, whose *V*_*1/2*_ at 25 °C was 3.9 mV more depolarized compared to 15 °C (p = 0.0052 in one-way ANOVA with Sidak’s multiple comparisons test) (Fig. 3, Table 1). Steady-state fast inactivation was complete in all tested channels except for Na_v_1.3 at 15 °C (Fig. 3). Similar to the activation process, the IV-curves became less steep at 15 °C, as indicated by increased slope factors (Table 1).

**Figure 3.**
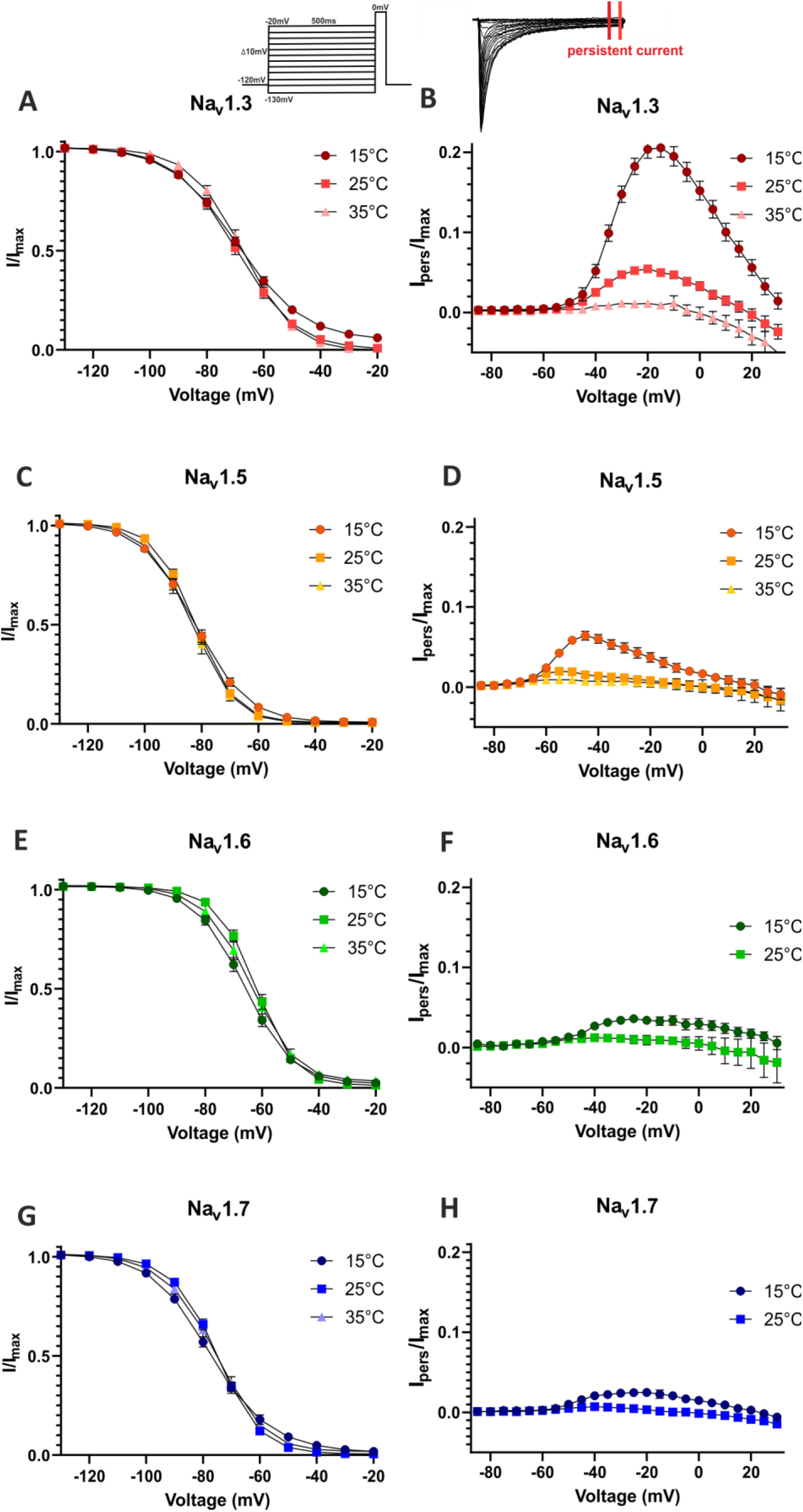
Temperature strongly modulates the kinetics, but not the voltage dependence of fast inactivation. Voltage dependence of steady-state fast inactivation at 15°C, 25°C and 35°C for **(A)** Na_v_1.3, **(C)** Na_v_1.5, **(E)** Na_v_1.6 and **(G)** Na_v_1.7. The voltage protocol is shown on top. **(B), (D)**, and **(H)** show persistent sodium current *I*_*pers*_ normalized to the maximum inward current *I*_*max*_ for the indicated subtypes and temperatures. All values are shown as mean ± 95 % confidence-interval.

The persistent current was enhanced with lowered temperature. This was the case for all investigated subtypes (Fig. 3B, D, F & H), but it was especially prominent for Na_v_1.3, where the maximum persistent current *I*_*pers*_ (as percentage of the maximum inward current) was about 4 times higher at 15 °C (21.0 ± 0.6 %) compared to 25 °C (5.2 ± 0.2 %), while at 35 °C nearly no persistent current was detectable (1.1 ± 0.1 %) (Fig. 3B). These results indicate that the voltage dependence of fast inactivation is only slightly affected, but the inactivation kinetics are strongly modulated by temperature.

We determined the inactivation time constant *τ* by a single exponential fit in order to further quantify the effect of temperature on inactivation kinetics. Rising temperature led to an acceleration of the inactivation kinetics with declining *τ*-values. Compared to 25 °C, *τ* was approx. 2.1 to 2.5 times larger at 15 °C and approx. 1.7 to 1.9 times smaller at 35 °C for Na_v_1.5, Na_v_1.6 and Na_v_1.7 at the voltage step to 0 mV (Table 1, Fig. 4D, G, J). For Na_v_1.3, lowering temperature from 35 °C to 25 °C led to a 2.2-fold and further cooling from 25 °C to 15 °C led to a 4-fold increase of *τ* (Fig. 4A). Compared to the *τ-*value of Na_v_1.7 at 15 °C (1.13 ms), that of Na_v_1.3 was 4.2 times larger (4.80 ms). This may indicate a special sensitivity of this channels inactivation kinetics towards cooling.

**Figure 4.**
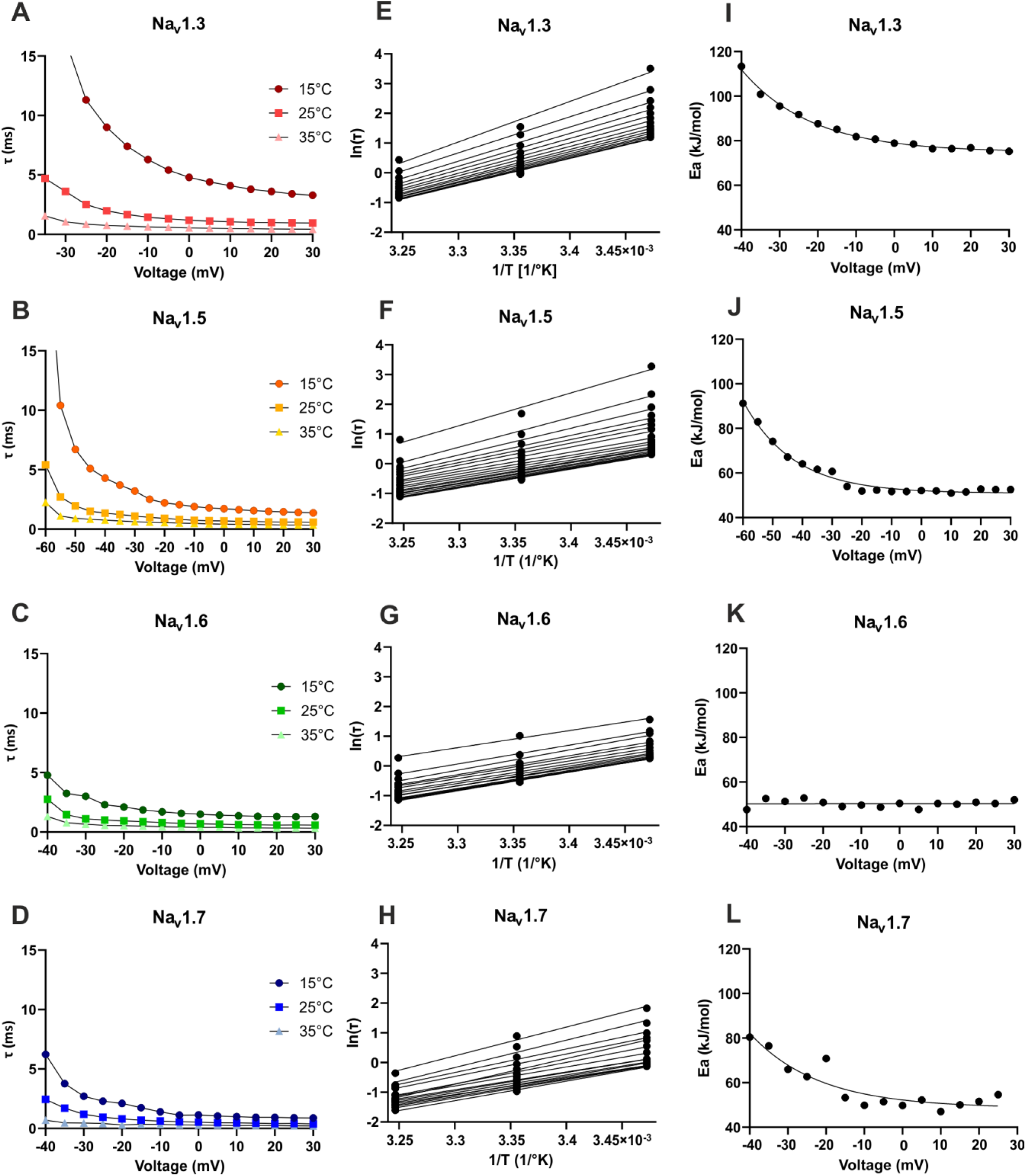
Kinetic and thermodynamic analysis of fast inactivation. **(A)-(D)** Inactivation time constant τ extracted from mono-exponential fits to the averaged traces at 15 °C, 25 °C and 35 °C. **(E)-(H)** Arrhenius plots calculated from inactivation time constant τ to determine the activation energy for the inactivation process at each voltage. **(I)-(L)** Summary of the Arrhenius analysis. Activation energy for inactivation plotted as a function of voltage.

An Arrhenius analysis was performed for potentials less negative than -40 mV (resp. -60 mV for Na_v_1.5). It revealed roughly flat plots only for Na_v_1.6, with an estimated *E*_*a*_ of ≈ 50 kJ/mol (Fig. 4G & K). Because *τ* = 1 / (α + β), this indicates that the forward rate constant α and the backward rate constant β from the open to the inactivated state have a similar *E*_*a*_ and τ ≈ 1/α can be assumed. For the other subtypes, *E*_*a*_ seems to depend on voltage, reflected in the curved Arrhenius plots (Fig. 4E, F & H) as well as the not linear progression of *E*_*a*_ plotted against voltage (Fig. 4I, J & L). The curves became flat only for potentials more positive than - 10 mV for Na_v_1.3 and Na_v_1.7 resp. -20 mV for Na_v_1.5, thus τ ≈ 1/α can be assumed only for this voltage range.

The averaged E_a_(α) for potentials less negative than -10 mV, was ≈ 52 kJ/mol for Na_v_1.5 and ≈ 51 kJ/mol for Na_v_1.7. For Na_v_1.3, it was 1.5 times larger, with ≈ 79 kJ/mol. This indicates, that for Na_v_1.3 more energy is needed to achieve fast inactivation, which may explain the strongly slowed inactivation of this subtype at 15 °C.

### The recovery from inactivation displays a high dependency on temperature

Because it is an important determinant of channel availability in high-frequency firing neurons we investigated the channels use-dependent current decay using the voltage protocol shown in Fig. 5A. Comparing the current of the first to the current of the last peak in a series of ten 5 ms depolarizations at 100 Hz, we observed a significantly stronger use-dependent current decay at 15 °C compared to 25 °C resp. 35 °C for all investigated subtypes (Fig. 5A).

**Figure 5.**
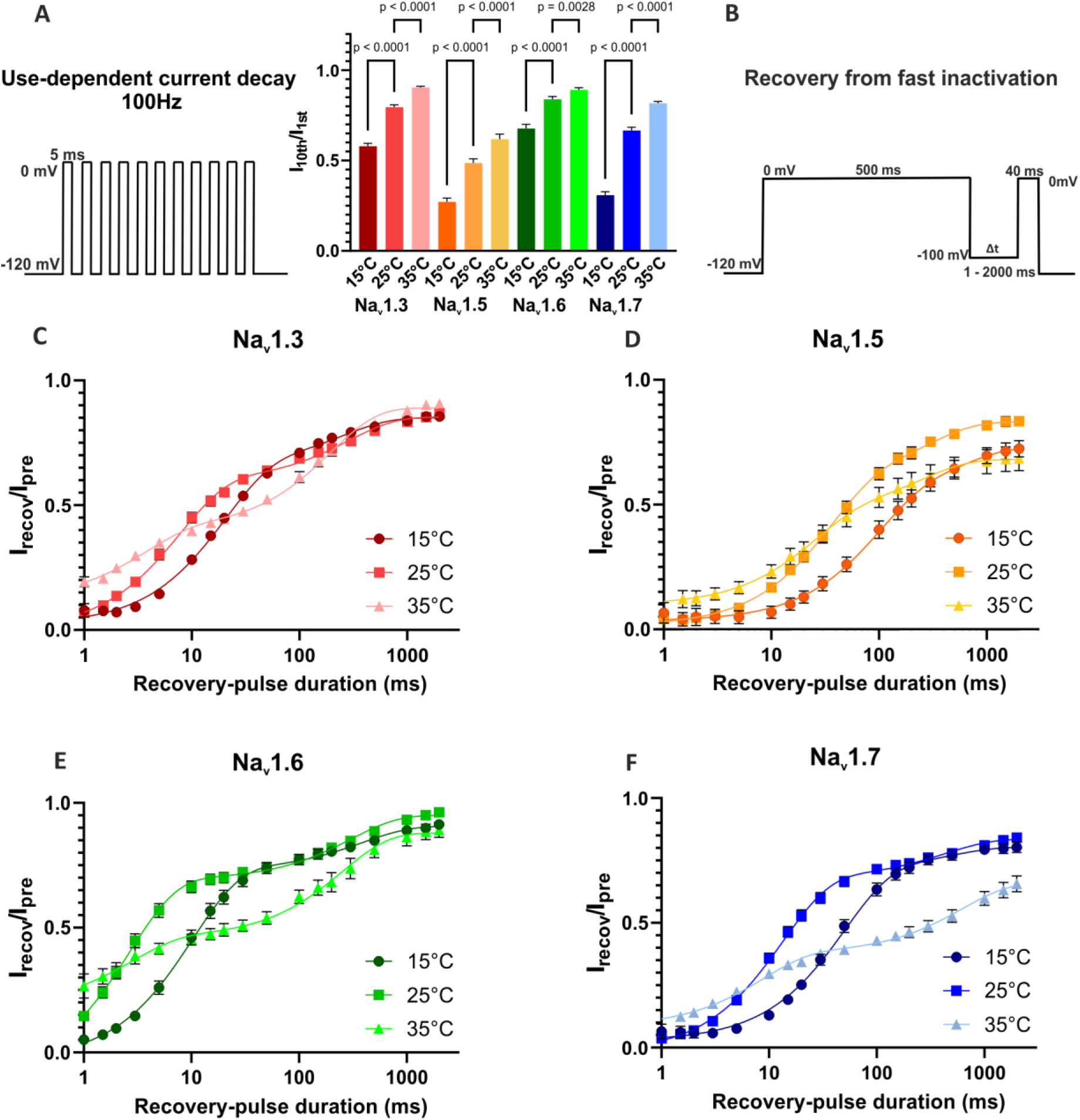
Temperature effects on use-dependent current decay and recovery from fast inactivation. **(A)** Use dependent current decay of Na_v_1.3, Na_v_1.5, Na_v_1.6 and Na_v_1.7 at 15°C, 25°C and 35°C at 100 Hz represented as the normalized current amplitude of the 10^th^ to the 1^st^ inward current. Voltage protocol shown on the left. All channel-subtypes show a statistically significant use-dependent current decay with decreasing temperature. **(B)-(E)** Recovery from fast inactivation at different temperatures. Normalized current amplitude as a function of recovery-pulse duration, voltage protocol shown on the top right. Data shown as mean ± 95% confidence-interval. One-way ANOVA with Sidak’s multiple comparisons test.

As recovery from fast inactivation has a large impact on use-dependent current decline, we used the voltage protocol shown in Fig. 5B to measure this gating characteristic. With warming from 15 °C to 25 °C we observed in general an acceleration of the channel’s recovery, expressed in the left shift of the curves (Fig. 5C – 5F) as well as decreased *τ*_*fast*_*-*values (Table 1). At 35 °C, lower *τ*_*fast*_-values indicate an even faster recovery compared to the one at 25 °C, but the proportion of fast recovering channels, *% τ*_*fast*_, decreased drastically, resulting in a flattening of the recovery curves and a much higher impact of the slow recovering process, represented by *τ*_*slow*_. With a pre-pulse duration of 500 ms, as used in this protocol, it is possible that a part of the channels is already slow inactivated. Our results suggest that, while the recovery from fast inactivation is accelerated at warmer, near physiological temperature, the onset of slow inactivation is enhanced at the same time.

### Warmth induced hyper- and cold induced hypoexcitability observed for the IEM-mutation Na_v_1.7/L823R

We showed that temperature has profound effects on Na_v_ gating of WT channels. Here, we investigated if cooling or warming affects biophysics of the disease related IEM causing mutation Na_v_1.7/L823R. The *V*_*1/2*_ of Na_v_1.7/L823R was significantly shifted to more hyperpolarized potentials with increasing temperature (Fig. 6F, Table 2). Compared to Na_v_1.7/WT, the mutation exhibited an approx. 10 mV hyperpolarizing shift of activation *V*_*1/2*_ at 25 °C and 35 °C and a 14.8 mV shift at 15 °C (p < 0.0001 in a Student’s t-test) (Fig. 6B & F, Table 2). This represents a fitting explanation for the patient’s phenotype with warm temperature triggering pain attacks. At 15 °C, the *V*_*1/2*_ of the mutation (*V*_*1/2*_ = - 37.4 ± 0.6 mV) was close to the *V*_*1/2*_ of the WT at 35 °C (*V*_*1/2*_ = - 35.4 ± 0.5 mV), suggesting that the voltage dependence of the channel’s activation is closer to the physiological (WT) conditions at colder temperature. Additionally, the L823R mutation renders the conduction curves less steep, with a slope factor *k* of 10.6 ± 0.3 mV at 15 °C, compared to the slope factor of 6.3 ± 0.2 mV for WT at 15 °C (Table 2). Comparing the time to peak values of WT and L823R mutation, we observed a slower activation at 25 °C (p < 0,005 in multiple unpaired t-tests), which was not detectable at 15 °C.

**Figure 6.**
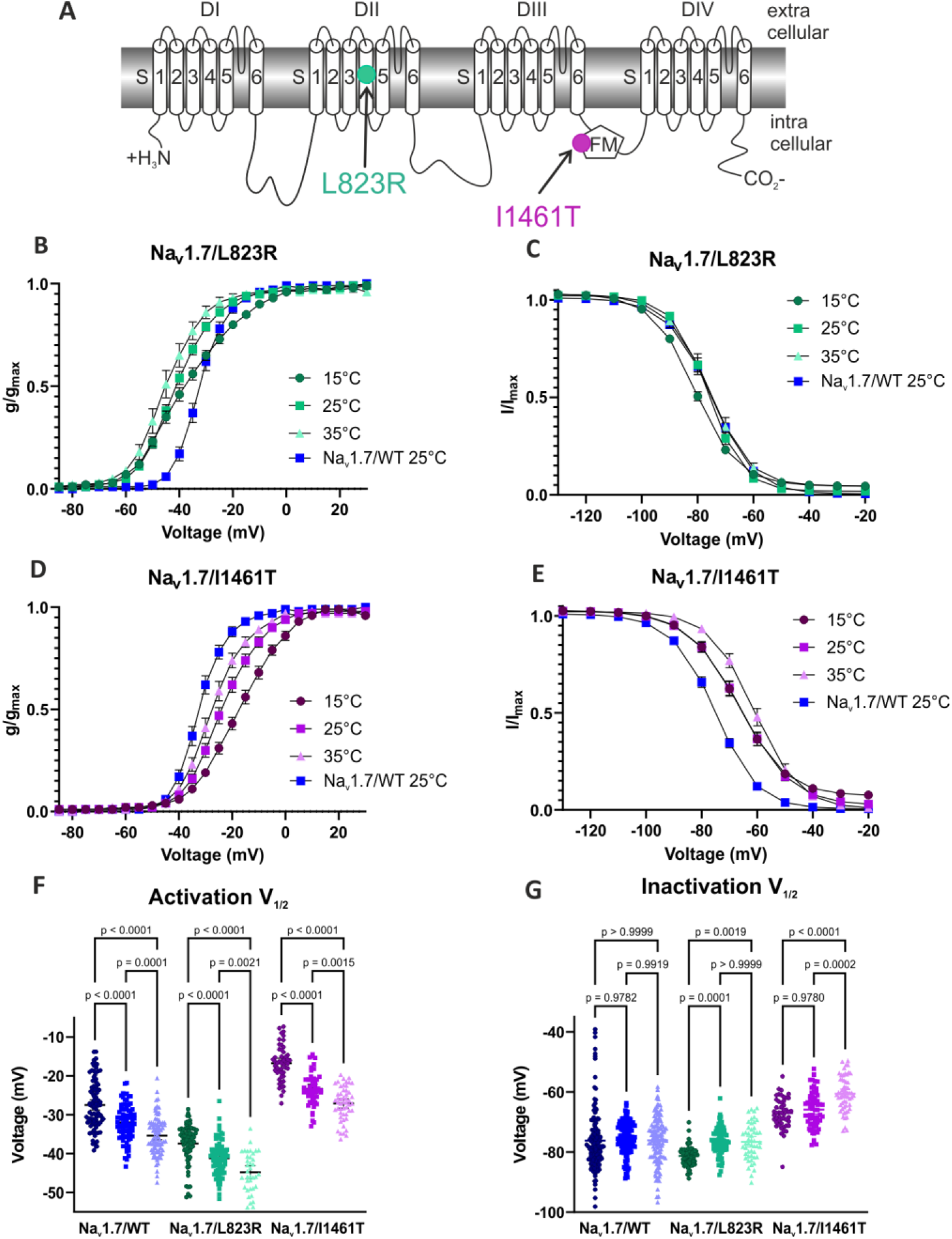
Voltage dependence of activation and steady-state fast inactivation of the IEM mutation Na_v_1.7/L823R and the PEPD mutation Na_v_1.7/I1461T compared to Na_v_1.7/I1461TWT. **(A)** 2D Scheme of the Na_v_1.7-channel showing the location of the affected amino acids for the mutation L823R (green symbol) and the mutation I1461T (purple symbol). **(B)** The G-V-dependence of Na_v_1.7/L823R is shifted to more hyperpolarized potentials compared to WT. **(C)** Steady-state fast inactivation of L823R is slightly shifted to more negative potentials at 15°C. **(D)** Activation and **(E)** steady-state fast inactivation are shifted to more depolarized potentials for Na_v_1.7/I1461T. **(F)** Values of half-maximal voltage dependence of activation and inactivation obtained from Boltzmann fits for individual traces. Data shown as mean ± 95% confidence-interval. One-way ANOVA with Sidak’s multiple comparisons test.

**Table 2.**
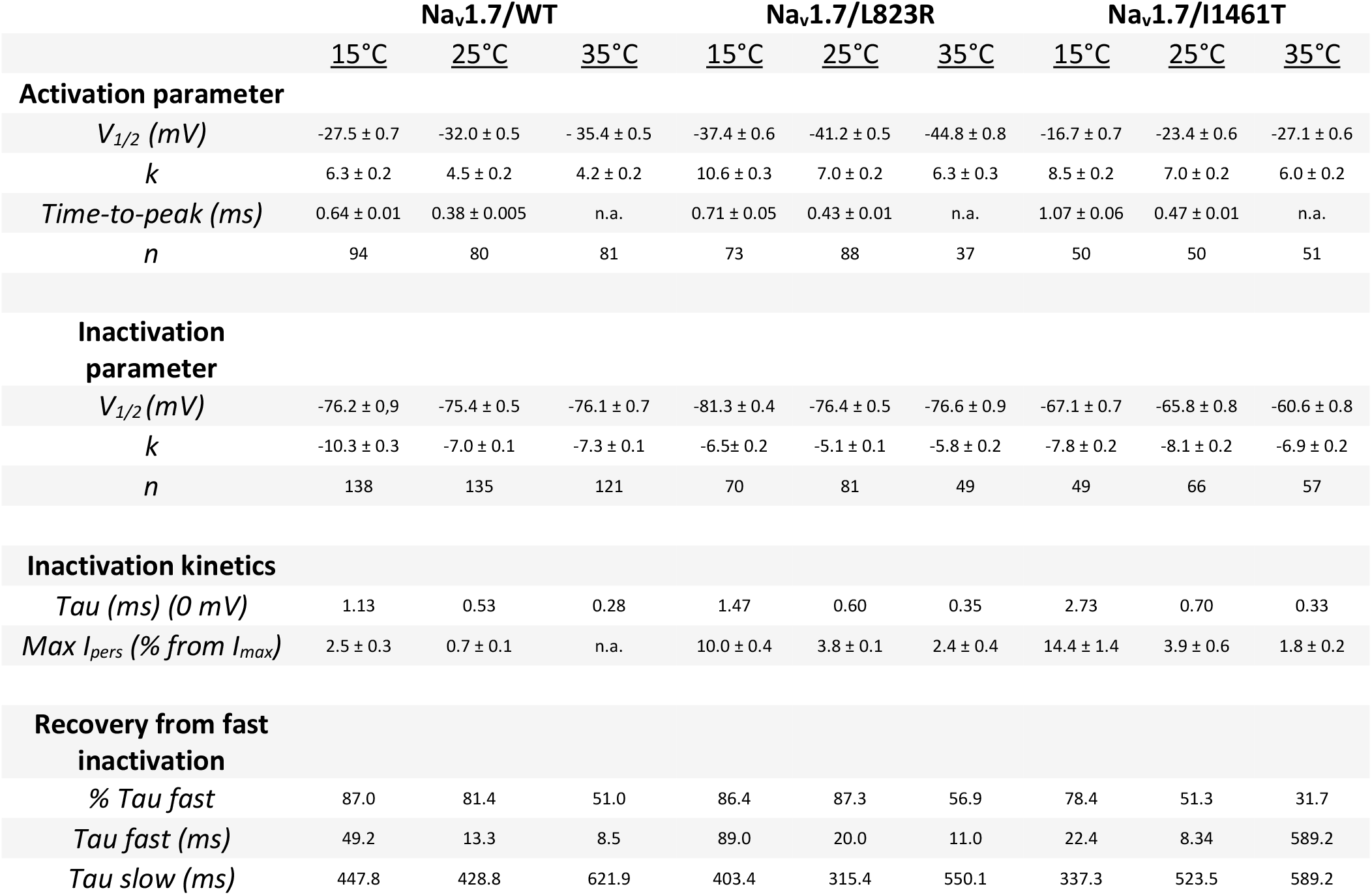
Summary of electrophysiological parameters determined for Na_v_1.7 and the two mutations Na_v_1.7/L823R and Na_v_1.7/I1461T (n.a.: not available)

In our experiments, we did not see any significant shift in the steady-state fast inactivation *V*_*1/2*_ of the mutation vs. WT at 35 °C or 25 °C (p = 0.6953 at 35 °C, p = 0.1452 at 25 °C in a Student’s t-test), but at 15 °C steady-state fast inactivation *V*_*1/2*_ was shifted by 4.8 mV to more hyperpolarized potentials for the mutation (p < 0.0001 in a Mann-Whitney test) (Fig. 6C & G, Table 2). This left shift in steady-state inactivation would also render neurons less excitable at 15 °C.

We noticed a 4-fold resp. 5.4-fold larger persistent current for the mutation at 15 °C and 25 °C (10.0 ± 0.4 % at 15 °C, 3.8 ± 0.1 % at 25 °C) than for the WT (2.5 ± 0.3 % at 15 °C, 0.7 ± 0.1 % at 25 °C) (Fig. 7A & B, Table 2). Increased leak current made accurate analysis of the persistent current difficult at 35 °C.

**Figure 7.**
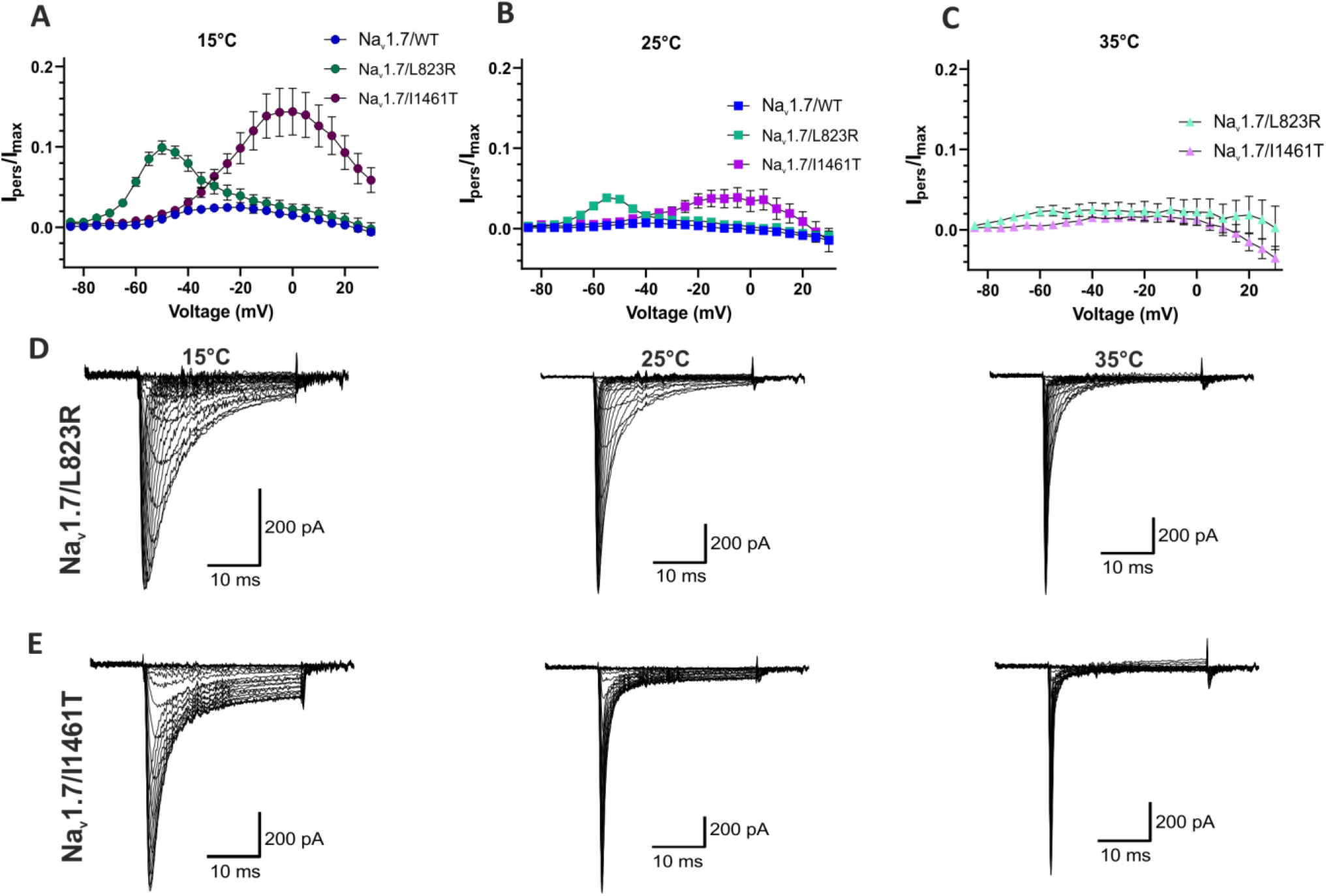
Cold temperature has stronger impact on both mutation’s inactivation kinetics than on Na_v_1.7WT. **(A)-(C)** Persistent current I_pers_ normalized to the maximal inward current I_max_. At **(A)** 15 °C and **(B)** 25 °C, the persistent current was strongly enhanced for both mutations compared to WT. Data shown as mean ± 95% confidence-interval. **(D)** Average of all included current traces for Na_v_1.7/L823R and **(E)** Na_v_1.7/I1461T.

The inactivation time constant *τ* of Na_v_1.7/L823R at 0 mV was approx. 1.2 times larger than for WT (Table 2, Fig. 8A). Thermodynamic analysis revealed Arrhenius plots (Fig. 8B) that were roughly parallel for the mutation for the voltage range above - 55 mV. With the averaged slope of this curves, *E*_*a*_ was estimated to be ≈ 55 kJ/mol, which is quite close to *E*_*a*_ ≈ 51 kJ/mol for Na_v_1.7/WT. In contrast to WT, no voltage dependence of *E*_*a*_ was observed (Fig. 8C).

**Figure 8.**
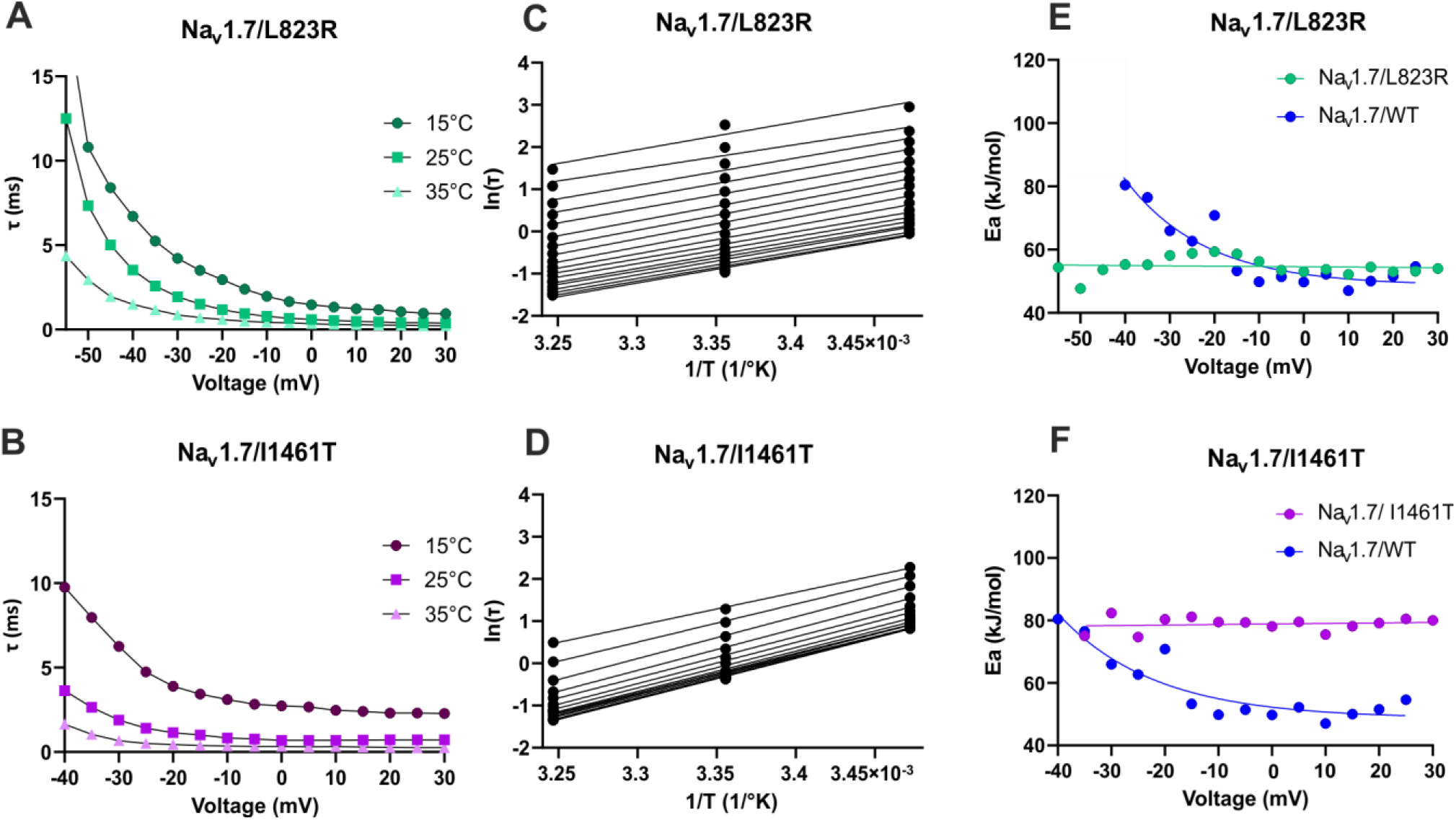
Kinetic and thermodynamic analysis of fast inactivation in Na_v_1.7 mutations. **(A)** Inactivation time constant *τ* at 15 °C, 25 °C and 35 °C for Na_v_1.7/L823R and **(B)** Na_v_1.7/I1461T. **(C)** and **(D)** Arrhenius plots calculated from *τ* to determine *E*_*a*_ for the inactivation process at each voltage. **(E)** Summery of the Arrhenius analysis with *E*_*a*_ of the inactivation process plotted as function of voltage, comparing Na_v_1.7WT with Na_v_1.7/L823R and **(F)** Na_v_1.7/I1461T. The average of all included current traces was used for this analysis.

Regarding the use-dependent current decay, we did neither observe any significant difference between L823R and WT for 50 Hz nor for 100 Hz at 15 °C, 25 °C and 35 °C (Fig. 9A). Testing for the recovery from fast inactivation, at 15 °C the recovering curve of the WT was slightly shifted towards shorter recovery times compared to the mutation (Fig. 9B), indicating a slower recovery process for the mutated channel. This was also reflected in the *τ*_*fast*_ value (45.63 ms for WT, 88.41 ms for L823R). At 25 °C, the recovery curves were almost overlapping with similar *τ*_*fast*_ values (Table 3, Fig. 9C) and at 35 °C the L823R channel recovered faster than the WT channel (Fig. 9C), but this result has to be treated with caution because the measurement of the mutation became unstable at 35 °C.

**Figure 9.**
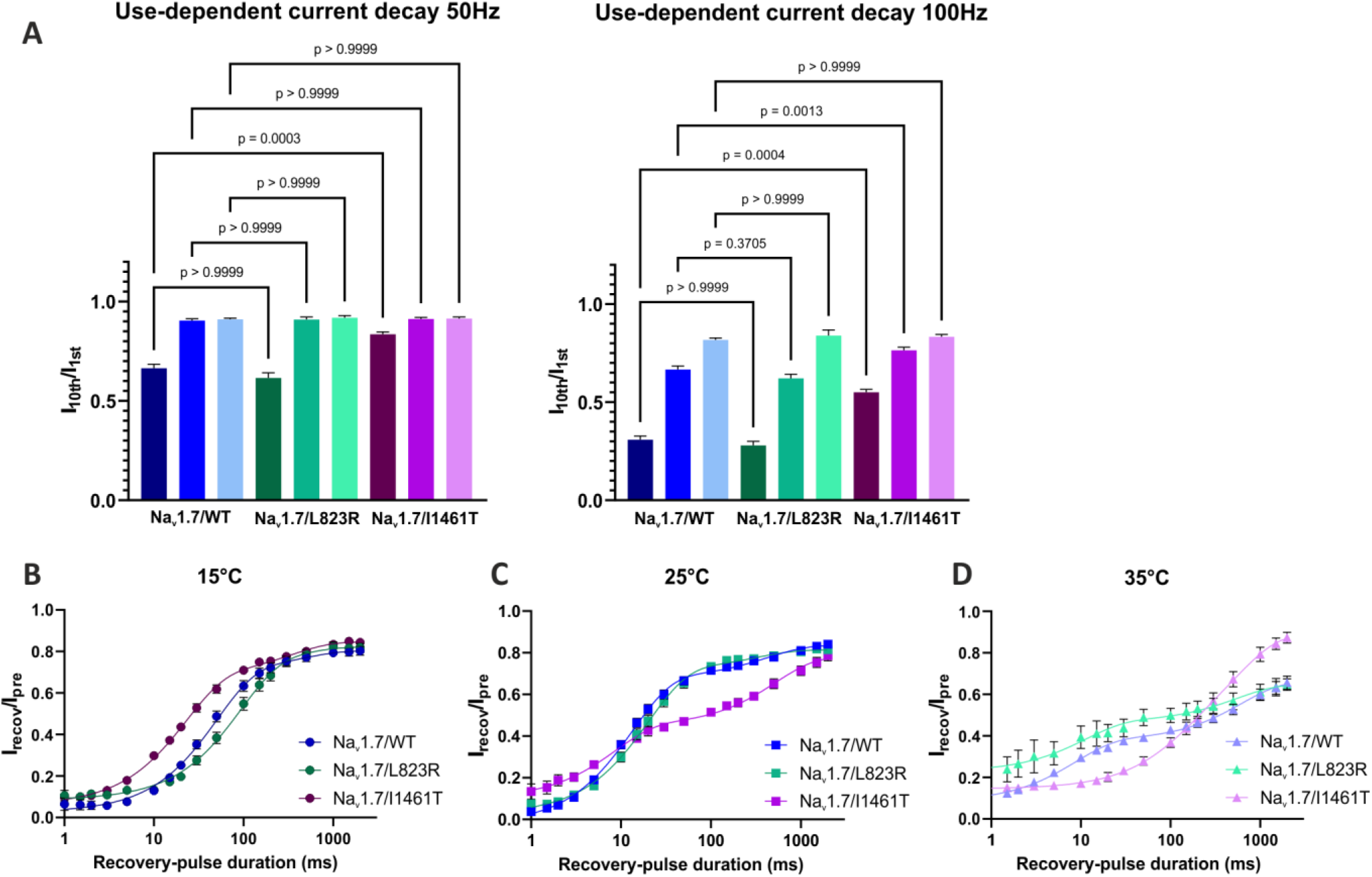
Na_v_1.7/I1461T shows a significant different behavior in use-dependent inactivation and recovery from inactivation compared to Na_v_1.7/WT and Na_v_1.7/L823R. **(A)** Use dependent current decay of Na_v_1.7/WT, Na_v_1.7/L823R and Na_v_1.7/I1461T at 15 °C, 25 °C and 35 °C represented as the normalized current amplitude of the 10^th^ to the 1^st^ inward current. At 15°C, Na_v_1.7/I1461T displayed significant smaller use-dependent inactivation compared to WT. **(B)-(D)** Recovery from fast inactivation, normalized current amplitude as a function of recovery-pulse duration. Data shown as mean ± 95% confidence-interval. Kruskal-Wallis test with Dunn’s multiple comparisons test. For recovery time-constants see Table 2.

It was described before that other mutations in Na_v_1.7 causing IEM increase ramp currents evoked by the application of slow depolarizing pulses (Han et al., 2007). Here, we applied three different ramps from a holding potential of -120 mV with a speed of 1.4 mV/ms, 2.5 mV/ms and 5 mV/ms. Fig. 10A shows example traces of ramp currents elicited by a 2.5 mV/ms ramp at 25 °C for Na_v_1.7/WT and both mutations. The ratio ramp current peak *I*_*ramp*_ to the maximum inward current *I*_*max*_ increased with decreasing temperature as well as with increasing ramp speed for all investigated subtypes (Fig. 10). Comparing Na_v_1.7/L823R with Na_v_1.7/WT, we observed a significant increase in ramp currents of the mutation at all tested temperatures (p < 0.0001 in Student’s t-tests). These results may indicate a higher sensitivity of the channels to slow subthreshold stimuli at temperatures between 15 °C and 35 °C.

**Figure 10.**
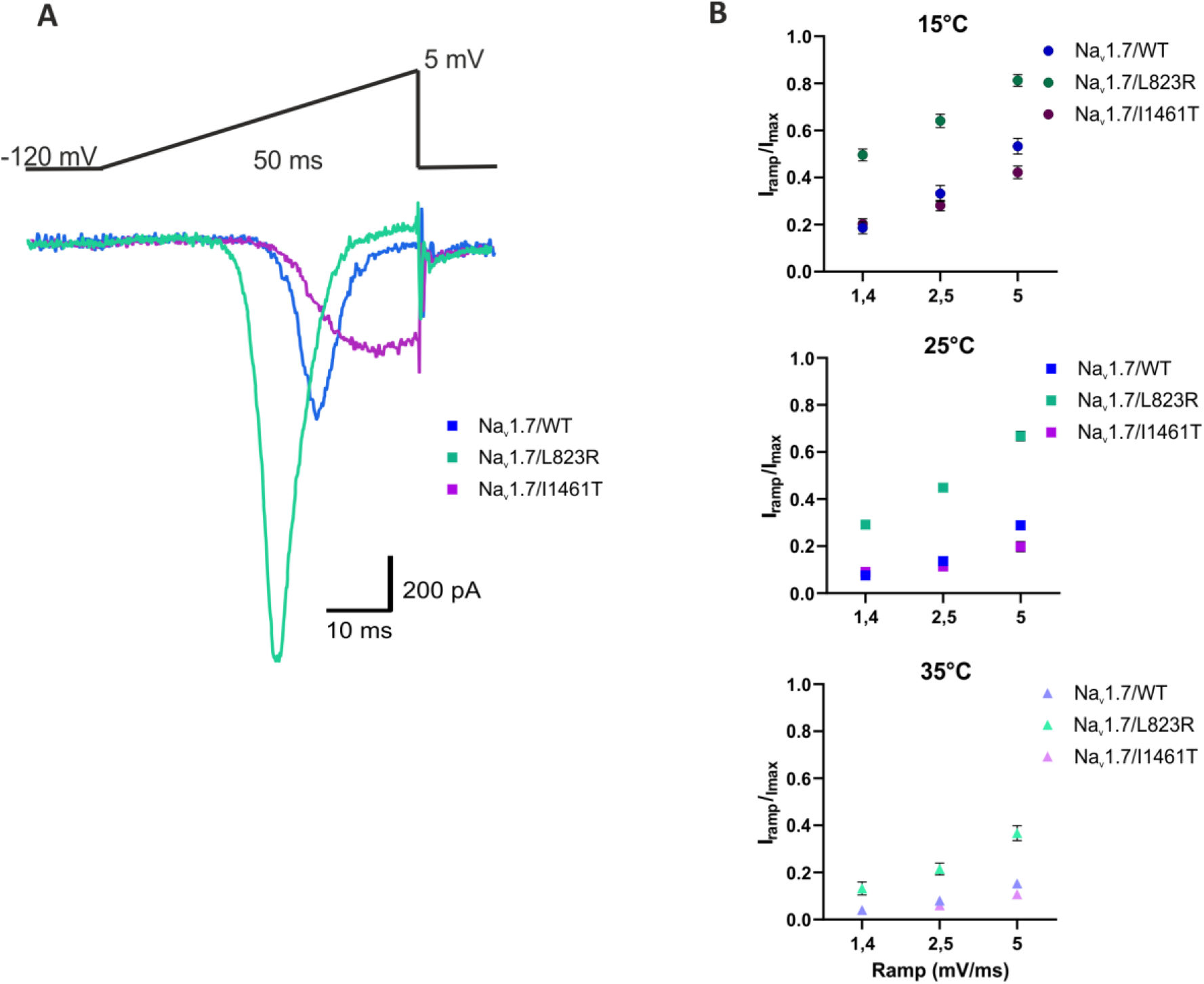
Na_v_1.7/L823R shows strongly enhanced ramp currents over the whole temperature range. **(A)** Example ramp-current elicited by 2.5 mV/ms ramps at 25 °C. Comparison of Na_v_1.7WT (blue), Na_v_1.7/L823R (green) and Na_v_1.7/I1461T (purple). **(B)** Ramp-current normalized to maximal inward current and plotted against ramp speed. The current increased with cooling and enhanced ramp-speed. Ramp current of Na_v_1.7/L823R was clearly stronger than the one of Na_v_1.7/WT and Na_v_1.7/I1461T. Data shown as mean ± 95% confidence-interval.

### Cooling induces an enhanced impaired inactivation for the PEPD mutation Na_v_1.7/I1461T

The PEPD mutation Na_v_1.7/I1461T, affecting the highly conserved inactivation motif in the DIII/IV-linker (Fig. 6A), was described before to induce changes in fast as well as in slow inactivation (Fertleman et al., 2006, Jarecki et al., 2008, Sheets et al., 2011). Here, we observed a shift to more depolarized potentials of steady-state fast inactivation compared to WT (Fig 6E & G). The *V*_*1/2*_ was approx. 10 mV less negative for the mutation at 15 °C and 25 °C (p < 0.0001 in Student’s t-tests) (Table 2). Warming to 35 °C shifted the IV-curve to even more depolarized potentials, an effect we did not observe in the steady-state fast inactivation of WT channels (Fig. 6G, Table 2).

For activation *V*_*1/2*_ we also observed shifts of 8.3 mV (35 °C), 8.6 mV (25 °C) and 11.8 mV (15 °C) to more depolarized potentials compared to WT (Fig. 6E, Table 2). Regarding the temperature dependence of activation, the mutation showed the same effect as WT with significant shifts in *V*_*1/2*_ to more hyperpolarized potentials in warmer conditions, from -16.7 ± 0.7 mV at 15 °C to -27.1 ± 0.6 mV at 35 °C (Fig. 6F). We observed a strong increase in persistent current for the mutation, with a normalized persistent current *I*_*pers*_ of 14.4 ± 1.4 % at 15 °C for Na_v_1.7/I1461T and only 2.5 ± 0.3 % for WT (Fig. 7A & B). Due to increasing leak current at 35 °C, the calculation of the persistent current was not precise at depolarized potentials (Fig. 7C & E).

The inactivation kinetics were slowed with decreasing temperature (Fig. 8B, Table 2). *τ-*values were, exemplary for the voltage step to 0 mV, 1.2 resp. 1.3 times larger at 35 °C and 25 °C compared to WT, but 1.9 times larger at 15 °C. Furthermore, *E*_*a*_ could be estimated for Na_v_1.7/I1461T for potentials less negative than -40 mV and showed no dependence on voltage in this range (Fig. 8 F). Averaged *E*_*a*_ of fast inactivation was ≈ 78 kJ/mol, a 1.6-fold increase compared to WT. The increased activation energy for fast inactivation may be explained by the impaired inactivation process of the mutation and causes probably the observed persistent current as well as the slowed inactivation.

Testing the channels use-dependent current decay at 50 Hz and 100 Hz, Na_v_1.7/I1461T displayed a stronger current decay with decreasing temperature, similar to Na_v_1.7/WT. However, at 15 °C it was significant smaller compared to WT for 50 Hz and at 100 Hz (Fig 9A).

A faster recovery from fast inactivation for the I1461T mutation was described before, which provides a possible explanation for this effect. Investigating the recovery from fast inactivation with a 500 ms pre-pulse to 0 mV revealed a speeding of the mutation’s recovery compared to WT at 15 °C, with *τ*_*fast*_ of 22.4 ms for I1461T and 45.6 ms for WT (Fig. 9B, Table2). Interestingly, at 25 °C the recovery behavior of the mutant was faster, as observed for the WT channel, but in addition the proportion of fast recovering channels (% *τ*_*fast*_) decreased from 78.4 % at 15 °C to 51.3 % at 25 °C (Table 2). Thus, the proportion of channels that recover slowly become more dominant. This resulted in a flattening of the curve (Fig. 9C) at 25 °C and 35 °C, which was only observed at 35 °C for the WT (Fig. 5F). Even though *τ*_*fast*_ is still smaller and for recovery-pulse durations up to 10 ms a higher proportion of mutant channels is recovering compared to WT, for longer recovery periods between 20 ms and 2000 ms the fraction of recovered channels is reduced for the mutation (Fig. 9C). This may indicate that slow inactivation is already occurring during the 500 ms pre-pulse at 25 °C. At 35 °C, the majority of the channels seem to mainly recover slowly, with a % *τ*_*fast*_ of only 31.7 % and a substantial proportion of Nav1.7/I1461T channels that just starts recovering when repolarized for periods longer than 50 ms (Fig. 9D).

Ramp currents of Na_v_1.7/I1461T were not significantly changed compared to WT at 15 °C, 25 °C and 35 °C for 1.4 mV/ms and 2.5 mV/ms ramps, while 5 mV/ms ramps were significantly smaller for the mutation (p < 0.0001 in a student’s t-test) (Fig. 10).

## Discussion

In this study, we investigated the effects of temperature on four different Na_v_ subtypes and two mutations of Na_v_1.7, which are linked to the inherited pain syndromes IEM and PEPD, under standardized conditions at 15 °C, 25 °C and 35 °C. We reveal a pronounced sensitivity of Na_v_1.3 fast inactivation kinetics to lowered temperature, resulting in a striking persistent current which may play a role in injury induced cold allodynia. Biophysical effects of pain-linked mutations in Na_v_1.7 were enhanced by warmth in IEM and by cooling in PEPD, which may explain the clinically observed specific triggers of these diseases.

### Temperature induced enhancement of Na_v_ activation

Here, we systematically investigated the temperature dependence of activation of Na_v_1.3, Na_v_1.5, Na_v_1.6, and Na_v_1.7 and observed a hyperpolarizing shift in the voltage dependence of activation with increasing temperature (Fig. 2 and 6). Due to the nature of our experimental setting, we can directly compare temperature effects on the channels and the large number of experiments due to high throughput patch clamp further increases the quality of our data. Temperature effects on Na_v_ activation was investigated before, mostly using manual patch clamp (Sarria et al., 2012, Thomas et al., 2009, Almog et al., 2022, Egri et al., 2012). However, reports on the direction of heat induced shifts in the voltage dependence of Na_v_ activation vary in literature depending on the experimental conditions, and some studies show no significant shift (Touska et al., 2018, Rosen, 2001, Ye et al., 2018) or even a shift of V_1/2_ to more depolarized potentials (Zimmermann et al., 2007) with increasing temperature. In our study, we used comparable conditions for all temperatures investigated and due to the high-throughput setting, our experiments have sufficient *n* to support our findings. The voltage dependence of Na_v_ activation is an important determinant for the excitability of the tissue it is expressed in. Thus, warmth induced shifts to more hyperpolarized potentials may explain for example increased neuronal excitability in febrile seizures (Thomas et al., 2009, Ye et al., 2018) or fever triggered arrhythmic events in normal hearts (Pasquie et al., 2004).

### V_1/2_ of steady-state fast inactivation is only slightly affected by temperature for WT Na_v_s

In this study, Na_v_1.3, Na_v_1.5 and Na_v_1.7 exhibited no significant shifts in the midpoint of fast inactivation induced by temperature and in case of Na_v_1.6 we observed only a slight shift to more hyperpolarized potentials comparing 15 °C to 25 °C (Fig. 3). For the voltage dependence of steady-state fast inactivation inconsistent temperature induced modulations were described in literature. Zimmermann et al. (2007), Ruff (1999), Ye et al. (2018) and Abdelsayed et al. (2013) reported significant hyperpolarizing shifts of fast inactivation *V*_*1/2*_ for Na_v_1.2, Na_v_1.4, Na_v_1.6, Na_v_1.7 and Na_v_1.8 with increasing temperature, while Egri et al. (2012) reported a depolarizing shift for Na_v_1.2. and Xiao et al. (2019) found no or only insignificant shifts for Na_v_1.2 and Na_v_1.8.

### Temperature intensifies effects of pain-linked mutations

Hyperexcitability induced by increased temperature is also reflected in the phenotype of patients suffering from IEM, who experience pain attacks that can be triggered by mild warmth (van Genderen et al., 1993, Albuquerque et al., 2011). Except for two, all known IEM mutations so far go along with hyperpolarizing shifts in the V_1/2_ of activation (Baker and Nassar, 2020, Choi et al., 2010, Eberhardt et al., 2014). Comparing the activation V_1/2_ of Na_v_1.7/L823R with Na_v_1.7/WT, we observed a hyperpolarizing shift of approx. 10 mV at all temperatures tested (Fig. 6B & F). The additional shift of V_1/2_ to more negative potentials, especially at 35 °C, may cause the neuronal hyperexcitability leading to pain sensation. Moreover, cooling leads to a depolarizing shift, bringing the midpoint of activation from L823R closer to the one of WT. Similar temperature induced effects were observed for the IEM mutation Na_v_1.7/L858F, which exhibited a depolarizing shift of activation V_1/2_ upon cooling to 16 °C (Han et al., 2007). This effect is in line with reports that pain can only be alleviated by cooling for most of IEM patients (van Genderen et al., 1993, Albuquerque et al., 2011, Tang et al., 2015). The physiological temperature of the skin is approx. 33 °C to 34 °C (Miland and Mercer, 2006), but already a one minute immersion into 15 °C cold water can reduce it to 17 °C (Dupuis, 1987) Thus, it is realistic that Na_v_1.7 channels that accumulate distally in nerve terminals of the skin are exposed to a large temperature range, and the skin can easily reach 15 °C or even cooler.

The steady-state fast inactivation *V*_*1/2*_ of the IEM linked Na_v_1.7/L823R was not shifted compared to WT at 25 °C and 35 °C, while at 15 °C a significant shift to more hyperpolarized potentials occurred (Fig. 6 C & G). This decreases the window current and thereby decreases the channel’s excitability at colder temperatures, what is in line with the clinical picture of IEM patients and the fact that cooling brings relief from pain. In patch clamp experiments at room temperature Lampert et al. (2009) also observed a shift of inactivation V_1/2_ of the mutant compared to WT. There, the search for a suitable explanation remained challenging because seen alone this would render the channel less excitable. This example points out that it is essential to perform electrophysiological investigations at different temperatures in order to understand the channel’s function in context and not to miss important effects.

In PEPD, patients also suffer from pain attacks, but in their case cold wind is reported as possible trigger factor (Fertleman et al., 2007). In contrast to the IEM mutation, a shift of activation of approx. 9mV to more depolarized potentials compared to WT was observed for the PEPD mutation Na_v_1.7/I1461T (Fig. 6D & F). In literature, no significant shift in activation V_1/2_ compared to WT was reported before for Na_v_1.7/I1461T, but PEPD causing mutations in the segment 4 (S4) - S5 linker of DIII, Na_v_1.7/V1298F and V1299F, as well as the mutation Na_v_1.7/T1464I showed shifts of 6.3 mV, 4.5 mV resp. 6.8 mV to more depolarized potentials (Fertleman et al., 2006, Jarecki et al., 2008). This is counterintuitive, because the shift would render cells expressing the channels less excitable when focusing on this gating mode only. We assume that the impaired inactivation that was observed for I1461T is sufficient to overcome the predicted reduction in excitability induced by the shift of activation and is also the crucial factor for the special sensitivity at colder temperature.

Ten of thirteen mutations causing PEPD that have been described so far are located in DIII and IV (Baker and Nassar, 2020). DIV plays an essential role in channels fast inactivation (McPhee et al., 1998, Capes et al., 2013, Goldschen-Ohm et al., 2013). Recently, Pan et al. (2018) proposed that rather than via direct occlusion, like it is described with the “hinged lid” mechanism, fast inactivation may occur because of an allosteric mechanism. The IFM motif binds to a hydrophobic pocket, thereby causing a movement of DIV S6 towards the ion permeation pathway leading to an occlusion. The hydrophobic cavity, to which the IFM motif binds, is formed by the S5 and S6 of DIII and IV and the DIII S4-5 linker (Pan et al. 2018, Shen et al 2019). For the PEPD mutation Na_v_1.7/A1632E, located in DIV S5, all-atom molecular dynamics simulations revealed that the glutamate side chain protrudes into the binding pocket and causes steric repulsion of the IFM motif. Thereby, it leads to impaired binding and thus impaired inactivation (Rühlmann et al., 2020). It is likely that changing the unpolar isoleucine of the IFM-motif in the Na_v_1.7/I1461T with a polar threonine results in a similar hydrophobic mismatch with impaired binding of the IFM into the hydrophobic cavity and a dysfunctional inactivation process.

We observed an approx. 10 mV depolarizing shift in steady-state fast inactivation of Na_v_1.7/I1461T at 15 °C and 25 °C compared to WT that was even larger at 35 °C. This increases the window current, rendering neurons carrying the mutation more excitable (Fig 6E & G). Our results confirm the depolarizing shift of fast inactivation that has previously been described for Na_v_1.7/I1461T (Fertleman et al., 2006, Jarecki et al., 2010, Jarecki et al., 2008, Jarecki et al., 2009) and for several other PEPD causing mutations at room temperature (Fertleman et al., 2006, Jarecki et al., 2008, Baker and Nassar, 2020, Dib-Hajj et al., 2008).

Temperature variations are also supposed to influence the Na_v_ gating kinetics. Here, we found an accelerated opening velocity, reflected in faster time-to-peak values observed with increasing temperature, similar to previous results (Touska et al., 2018, Almog et al., 2022). Faster opening could also promote increased excitability at elevated temperatures. Even though the L823R mutation induces an additional positive charge in the voltage sensor of DII, the activation kinetics were slightly slower than for WT. This effect was observed before and it was stated that it might be induced by an unusual rearrangement during the opening process caused by the extra arginine, with DII moving prior to DIII, and thus increasing the time needed for opening (Lampert et al., 2009).

### Subtype-specific modulation of inactivation kinetics

Our study revealed that Na_v_ inactivation kinetics are slowed with decreasing temperature resulting in a larger value of the inactivation time constant, similar to what has been reported by several others before (Zimmermann et al., 2007, Thomas et al., 2009, Ke et al., 2017, Sittl et al., 2012, Egri et al., 2012, Touska et al., 2018). The comparison of different subtypes, which was possible under standardized conditions in this study, showed that, even though cooling has the same overall effect on all investigated Na_v_s, the extent of the effects differs strongly among subtypes, especially when focusing on the persistent current (Fig. 4). Compared to Na_v_1.7/WT at 15 °C, Na_v_1.3 had a more than 8 times larger persistent current and a 4 times slower inactivation and the high persistent current nearly diminished at 35 °C. Differences in inactivation kinetics among the other subtypes were considerably smaller. This suggests Na_v_1.3 might have an important role in mediating cold or cold related sensations.

Interestingly, we observed enhanced persistent current at 15 °C and 25 °C also for Na_v_1.7/L823R compared to WT (Fig. 7). Persistent current of L823R was mainly detected in the voltage range between -70 mV to -30 mV, where not all channels are activated yet. Persistent currents in this voltage range might be related to the impaired balance of activation and inactivation, induced by the shift of activation to more hyperpolarized potentials, while steady-state fast inactivation V_1/2_ is at the same time nearly unaffected, but strongly slowed down by lowered temperature. In contrast, the large persistent current of Na_v_1.7/I1461T at 15 °C is probably caused by an impaired inactivation mechanism.

### Thermodynamic analysis

With the Arrhenius analysis of the inactivation time constant *τ*, an estimation of the activation energy *E*_*a*_ for the transition from the open to the inactivated state was possible. Averaged values of *E*_*a*_ at potentials more depolarized than -10 mV were between 50 to 55 kJ/mol for Na_v_1.5, Na_v_1.6, Na_v_1.7 and Na_v_1.7/L823R. Na_v_1.3 and Na_v_1.7/I1461T displayed a clearly higher *E*_*a*_, with ≈ 79 kJ/mol resp. ≈ 78 kJ/mol (Fig. 4 & 8). This may explain the drastically slowed inactivation and the large persistent current observed at 15 °C for these two channels. More energy is needed to inactivate them properly, and it is lacking at colder temperature. Interestingly, Na_v_1.6 as well as both mutations showed no voltage dependence of *E*_*a*_, while the other subtypes did so. This indicates that the forward rate constant α and the backward rate constant β have the same *E*_*a*_ over the whole voltage range and not only at depolarized potentials.

### Enhanced excitability at cold temperature as a result of persistent current and a potential role of resurgent current

A slowed or destabilized inactivation and large persistent current may lead to increased resurgent current (Hampl et al., 2016). Current clamp experiments as well as computer models have linked resurgent currents to neuronal hyperexcitability (Jarecki et al., 2010, Sittl et al., 2012, Xiao et al., 2019). An increase in resurgent current has been observed for Na_v_1.7/I1461T and other PEPD mutations as well as other Na_v_ mutations slowing fast inactivation kinetics and a correlation between the current decay time constant and resurgent current has been revealed (Jarecki et al., 2010, Theile et al., 2011, Sittl et al., 2012, Xiao et al., 2019, Hampl et al., 2016, Eberhardt et al., 2014). According to this, we assume a possible increase in resurgent current not only for the I1461T mutation, but also for Na_v_1.3 at 15 °C. Experiments with Oxaliplatin treated nerve fibers revealed Na_v_1.6 mediated enhanced persistent and resurgent currents that were linked to cold aggravated neuropathy (Sittl et al., 2012).

Taken together, a strongly slowed inactivation and thus enhanced persistent current can be induced by either a mutation (like for Na_v_1.7/I1461T) or cooling temperature (like for Na_v_1.3) or both, making it a potentially general mechanistic basis for cold aggravated symptoms and a possible explanation for the cold induced hyperexcitability as observed in patients suffering from PEPD. Cold sensitive mutations in Na_v_1.4 causing PMC are also supporting this hypothesis. PMC induces muscle stiffness in response to lowered ambient temperature. Several PMC mutations were shown to shift V_1/2_ of fast inactivation to more depolarized potentials, slow the current decay time constant and accelerate the recovery from fast inactivation, similar to PEPD mutations (Bouhours et al., 2004, Farinato et al., 2019, Palmio et al., 2017, Ke et al., 2017).

### Na_v_1.3 overexpression may be crucial for cold allodynia after nerve injury

Na_v_1.3 exhibited a special sensitivity towards cooling compared to the other investigated subtypes, with exclusively large persistent currents and slow inactivation kinetics (Fig. 3D & E). It is primarily expressed during the development of the nervous system, but it is expressed in only small amounts in adult DRG neurons (Felts et al., 1997, Chang et al., 2018). It was shown, that Na_v_1.3 overexpression occurs in rat DRG neurons after peripheral axotomy (Dib-Hajj et al., 1996) or spinal cord injury (SCI) (Hains et al., 2003, Lampert et al., 2006), leading to hyperexcitability of DRG nociceptive neurons and pain related behavior due to rapidly repriming TTX-sensitive currents. Dorsal horn neurons of rats with SCI displayed enhanced persistent and ramp currents, supporting hyperexcitability (Lampert et al., 2006). Lindia et al. (2005) showed a higher sensitivity towards cold stimuli in rats after SCI compared to unaffected controls. Our results indicate that cold sensitivity after SCI may be explained by the overexpression of Na_v_1.3, whose inactivation kinetics are slowed by cooling, thus potentially leading to enhanced resurgent currents and hyperexcitability at low temperatures. Even though we carried out the experiments in HEK expressing rNa_v_1.3 - not neuronal cells - and differences in the inactivation kinetics have been described for rNa_v_1.3 compared to hNa_v_1.3 (Tan and Soderlund, 2009), our results suggest that Na_v_1.3 overexpression may be a determining factor in the development of cold allodynia. Further investigation on the role of Na_v_1.3 in neuropathic pain is needed, such as exploring its potential as a candidate as a therapeutic target in the therapy of cold allodynia.

### The recovery from fast and the onset of slow inactivation is accelerated 35 °C

For all of the investigated Na_v_-WT channels as well as the Na_v_1.7/L823R mutation, our investigations revealed an acceleration of the recovery from inactivation when increasing the temperature from 15 °C to 25 °C. The rate of fast inactivation recovery increases even more at 35 °C, but only for recovery times of 3 ms to 12 ms (depending on the subtype) and the fast recovering proportion of channels (% *τ*_*fast*_) decreases (Fig. 5 & 9, Table 1 & 2). With longer recovery periods, the recovery rate was smaller at 35 °C compared to 25 °C, suggesting that more channels are recovering slowly from inactivation with increasing temperature. We assume that with the 500 ms pre-pulse, which we used in our experiments, a slow inactivation component was detected in the investigation of recovery from fast inactivation at 35 °C, reflected in the unexpected curve shapes and the high percentage of channels slowly recovering. This is consistent with the observation of a faster onset of slow inactivation with elevated temperature reported in the literature (Ke et al., 2017, Egri et al., 2012).

Taken together, our results suggest that with increasing temperature the recovery from fast inactivation is accelerated, but at the same time, the entry into slow inactivation seems to be enhanced. This may result in a higher availability of channels in high-frequency firing neurons at 35 °C compared to 25 °C and 15 °C, but it would also protect Na_v_s from excessive firing resulting in hyperexcitability.

At 15 °C Na_v_1.7/I1461T was recovering more quickly and use-dependent current decay was reduced compared to WT (Fig. 9A). Thus, neurons carrying the mutation are more likely to show high frequency firing at colder temperature than neurons expressing the WT channel, an effect that was described at room temperature before (Jarecki et al., 2008, Sheets et al., 2011). Compared to WT the I1461T mutation started to show a significant slower recovery from fast inactivation for recovery-pulse durations longer than 10 ms, resulting in a flattening of the curve already at 25 °C, while for the WT this effect was only observed at 35 °C (Fig. 9B). At this temperature, the mutated Na_v_s recovered mainly slowly, with a %*τ*_*fast*_ of only 37 %, indicating that the proportion of channels recovering slowly was even more enhanced at this temperature. Even though Jarecki et al. (2008) described a decrease in the voltage dependent transition into slow inactivated states for Na_v_1.7/I1461T and Sheets et al. (2011) observed no differences in the development of slow inactivation for WT and mutation, our observations suggest that this might be changed with increasing the temperature to 25 °C or more. Further studies on the onset as well as recovery from fast inactivation are for sure necessary to get deeper insights into the mechanisms and temperature induced effects. However, in combination with the results of the inactivation time constant and the enhanced persistent currents at 15 °C, these results reinforce the special sensitivity of the I1461T mutation towards temperature changes in both directions, warming and cooling, and gives evidence that neuronal hyperexcitability causing pain in the context of PEPD may be inducible by cooling.

### Putative role of the β1-subunit

The impact of β-subunits on the gating of Na_v_s should not be neglected in the discussion of temperature induced changes of excitability. Next to the effect of β-subunits on the voltage dependence of gating states, resurgent currents (Grieco et al., 2005), stabilization against mechanical stimuli (Körner et al., 2018), and a possible contribution to intercellular communication and recruitment of Na_v_s (Malhotra et al., 2000), Egri et al. (2012) revealed a thermoprotective role of the β1-subunit. Temperature induced changes to excitability are modulated by the expression of the β1-subunit with the general trend that at elevated temperature channels without β1 spend less time in the inactivated state. Thus, mutations in β1 can also lead to hyperexcitability at increased temperature, like it was shown for the mutation β1(C121W), causing epilepsy with febrile seizures plus, a pediatric febrile seizure syndrome.

### Future investigations and outlook

Automated patch-clamp systems can nowadays perform high-throughput electrophysiological experiments and can at the same time adjust the temperature in a range from 10 °C to 43 °C. With this, the technical challenges associated with performing electrophysiological experiments at elevated temperatures with manual patch clamp techniques can be overcome. At the same time, the amount of obtained data increases by a multiple. The quality of the data is comparable to those of manual patch clamp, with only slightly higher series resistances (compare methods) and seal resistances above 1 GΩ, that were maintained over long periods during the experiments. At 35 °C we sometimes experienced trouble with leak currents that weren’t subtracted properly (Fig. 1) and overshooting large sodium currents to more than 10 nA led to some inaccuracies in series resistance compensations. With stable cell lines, on the other hand, we were able to achieve success-rates of more than 80 %, even in recordings at 35 °C.

For drug testing and safety pharmacology it is crucial to gain deeper insight into gating mechanisms at physiological temperature of different ion channels. The state dependence of drug binding is likely to be coupled with the occupancy of different conformational states of the channel (Sheets et al., 2011) and this is in turn depending on temperature. E-4031, an experimental class III antiarrhythmic drug, was for example fivefold more potent in blocking hERG channels at 35 °C than at 22 °C (Davie et al., 2004). Thus, to investigate accurate potencies of drugs that modulate ion channels, it would be preferable to perform these studies at 37 °C. Detailed electrophysiological investigation of Na_v_s at different temperatures is also essential to understand the properties and to gain a deeper insight into the complex gating mechanisms of Na_v_s at physiological temperature, otherwise important effects might be missed. Of course, in the physiological context, the excitability of a tissue does not only depend on one single channel, but the interplay between many different channels. Furthermore, passive membrane properties have shown to be temperature sensitive and alter excitability (Touska et al., 2018, Volgushev et al., 2000). However, biologically detailed neuronal simulation models contain still crude assumptions about the kinetics of Na_v_s and many poorly constrained parameters, like Q10 values, that should only be used carefully because of their temperature and voltage dependence (Almog and Korngreen, 2016).

Thus, it is of enormous importance to collect experimental data of different ion channels also at physiological temperatures, to improve the reliability of action potential simulations and neuronal models, draw (patho-) physiological relevant conclusions and thus understand Na_v_s in their physiological context.

## Acknowledgements

We thank Jannis Körner for thoughtful discussions of the patch clamp recordings and Lennart Müller for technical support.

This work was supported by the 2020 SyncroPatch384i award by Nanion Technologies GmbH, Munich, Germany to AL and RH. This study was funded by grants from the Interdisciplinary Centre for Clinical Research within the faculty of Medicine at the RWTH Aachen University [IZKF TN1-1/IA 532001 and TN1-5/IA 532005] andby the Deutsche Forschungsgemeinschaft (German Research Foundation LA 2740/3–1, 363055819/GRK2415 Mechanobiology of 3D epithelial tissues (ME3T); 368482240/GRK2416, MultiSenses-MultiScales).

Angelika Lampert has a consultancy agreement and a research contract with Grünenthal. The authors declare no competing financial interests.

## Authors contributions

Sophia Kriegeskorte, Raya Bott, Martin Hampl, Ralf Hausmann and Angelika Lampert planned and Sophia Kriegeskorte, Raya Bott and Martin Hampl performed the patch clamp experiments. Sophia Kriegeskorte analyzed the patch clamp experiments, designed the figures and wrote the manuscript. Alon Korngreen performed the thermodynamic analysis. All authors discussed the results and the final manuscript. Angelika Lampert and Ralf Hausmann conceived the study, allocated funding and supervised the project.

