## Supplemental Table 1 for "Cold and warmth intensify pain-linked sodium channel gating effects and persistent currents"

### Supplemental material

***Table S1.*** *Summary of cell-culture media and supplements which were used for cultivation of HEK293 rNa$_v$1.3, hNa$_v$1.5, mNa$_v$1.6, hNa$_v$1.7/WT, hNa$_v$1.7/L823R, and hNa$_v$1.7/I1461T*

| Stable cell-line | Cell-culture media and supplements |
|---|---|
| **HEK293 rNa$_v$1.3** (Cummins et al., 2001) | Dulbecco´s modified Eagle medium (DMEM) with 4,5 % Glucose and L-Glutamine (Thermo Fisher Scientific, Waltham, Massachusetts, USA)<br>10 % fetal bovine serum (FBS; Sigma-Aldrich, St. Louis, Missouri, USA)<br>0,5 mg/ml Genetecin (G418; Carl Roth GmbH+Co.KG, Karlsruhe, Germany) |
| **HEK293 hNa$_v$1.5** (Eberhardt et al., 2015) | Dulbecco´s modified Eagle medium F-12 (DMEM/F1-2; Thermo Fisher Scientific, Waltham, Massachusetts, USA), 10 % FBS, 100 µg/ml Zeocin (Invivogen, San Diego, California, USA) |
| **HEK293 mNa$_v$1.6** (Herzog et al., 2003, Laezza et al., 2009) | DMEM/F-12, 10 % FBS, 0,5 mg/ml G418 |
| **HEK293 hNa$_v$1.7** (From Anaxon AG, Berne, Switzerland) | Ham´s F-12 Nutrient Mix, GlutaMAX$^{TM}$ Supplement (Thermo Fisher Scientific, Waltham, Massachusetts, USA), 9 % FBS, 1 mM Sodium Pyruvate (Thermo Fisher Scientific, Waltham, Massachusetts, USA), 150 µg/ml Hygromycin B (Carl Roth GmbH+Co.KG, Karlsruhe, Germany) |
| **HEK293 hNa$_v$1.7/L823R** (Merck KGaA, Darmstadt, Germany) | DMEM, 10 % FBS, 0,1 mM MEM Non-Essential Amino Acids Solution (NEAA; Thermo Fisher Scientific, Waltham, Massachusetts, USA), 25 mM HEPES (Carl Roth GmbH+Co.KG, Karlsruhe, Germany), 1 mg/ml G418, 5 µg/ml Blasticidin (Thermo Fisher Scientific, Waltham, Massachusetts, USA)<br>(1 µg/ml Doxycycline (Sigma-Aldrich, St. Louis, Missouri, USA) added 24 hours prior to the experiment) |
| **HEK293 hNa$_v$1.7/I1461T** (Merck KGaA, Darmstadt, Germany) | DMEM, 10% FBS, 0,1 mM NEAA, 25 mM HEPES, 10 µg/ml Blasticidin |
